# Analysis of Gene Expression from Systemic Lupus Erythematosus Synovium Reveals a Profile of Activated Immune Cells and Inflammatory Pathways

**DOI:** 10.1101/2020.06.19.123307

**Authors:** Erika L. Hubbard, Michelle D. Catalina, Sarah Heuer, Prathyusha Bachali, Robert Robl, Nicholas S. Geraci, Amrie C. Grammer, Peter E. Lipsky

## Abstract

Arthritis is a common manifestation of systemic lupus erythematosus (SLE) yet understanding of the underlying pathogenic mechanisms remains incomplete. We, therefore, interrogated gene expression profiles of SLE synovium to gain insight into the nature of lupus arthritis (LA), using osteoarthritis (OA) and rheumatoid arthritis (RA) as comparators. Knee synovia from SLE, OA, and RA patients were analyzed for differentially expressed genes (DEGs) and also by Weighted Gene Co-expression Network Analysis (WGCNA) to identify modules of highly co-expressed genes. Genes upregulated and/or co-expressed in LA revealed numerous immune/inflammatory cells dominated by a myeloid phenotype, whereas OA was characteristic of fibroblasts and RA of T- and B-cells. Upstream regulator analysis identified *CD40L* and inflammatory cytokines as drivers of the LA gene expression profile. Genes governing trafficking of immune cells into the synovium by chemokines were identified, but not *in situ* generation of germinal centers. GSVA confirmed activation of specific myeloid and lymphoid cell types in LA. Numerous therapies were predicted to target LA, including TNF, NFκB, MAPK, and CDK inhibitors. Detailed gene expression analysis identified a unique pattern of cellular components and physiologic pathways operative in LA, as well as drugs potentially able to target this common manifestation of SLE.

## INTRODUCTION

Systemic lupus erythematosus (SLE) is a complex autoimmune disease in which loss of self-tolerance gives rise to pathogenic autoantibodies causing widespread inflammation and tissue damage [1]. Lupus arthritis (LA) is a common manifestation of SLE with 65 to 95% of lupus patients reporting joint involvement during the course of their disease [2].

Despite the high frequency of LA, an understanding of the underlying pathogenic mechanisms remains incomplete. Cytokines, such as IL-6, and anti-dsDNA autoantibodies are thought to play a role [3–5]. Other autoantibodies including anti-ribonucleoprotein, anti-histone, and anti-proliferating cell nuclear antigen have been implicated in LA along with evidence of increased C-reactive protein (CRP) and erythrocyte sedimentation rate (ESR) [4, 6, 7].

The lack of a better understanding of the nature of LA relates to the difficulty of obtaining tissue samples and the absence of relevant and reliable animal models. Despite this, in most recent clinical trials of potential lupus therapies, arthritis is a principal manifestation and the success of a tested therapy can depend on its ability to suppress synovial inflammation. Therefore, it is essential to understand more about the pathogenic mechanisms operative in LA.

One way to evaluate the pathologic processes involved in LA is to analyze gene expression profiles in the affected synovium. Previous work analyzed global gene expression profiles and histology of SLE, rheumatoid arthritis (RA), and osteoarthritis (OA) synovium [8–9] to begin to elucidate the inflammatory mechanisms in each disease and focused on the type 1 interferon pathway in LA. Here, we expand upon these studies by applying contemporary bioinformatic techniques to assess the only gene expression data set available to gain additional insight into the pathogenesis of LA. Using a multipronged, bioinformatic and systems biology approach, we provide an expanded view of SLE synovitis that might serve as the basis to identify new targeted therapies.

## RESULTS

### Bioinformatic and Pathway Analysis of LA and OA Gene Expression

Differential expression (DE) analysis demonstrated 6,496 differentially expressed genes (DEGs) in LA versus OA (Fig. 1a), of which 2,477 transcripts were upregulated whereas 4,019 transcripts were downregulated. The upregulated DEGs included 243 immune cell-specific transcripts (odds ratio of 2.84, p<2.2e-16, Fisher’s Exact Test), indicating a significant immune/inflammatory cell infiltrate. There was considerable enrichment of T-cell, B-cell, plasma-cell, and myeloid-cell transcripts among the upregulated DEGs (Fig. 1b), whereas fibroblast-associated genes were increased in OA. Immune signaling and immune cell surface markers, as well as the interferon transcriptional program, pattern recognition receptors (PRRs), and MHC Class I and II were enriched in LA, along with intracellular signaling, apoptosis, the proteasome, and processing/packaging material inside cells.

**Figure 1.**
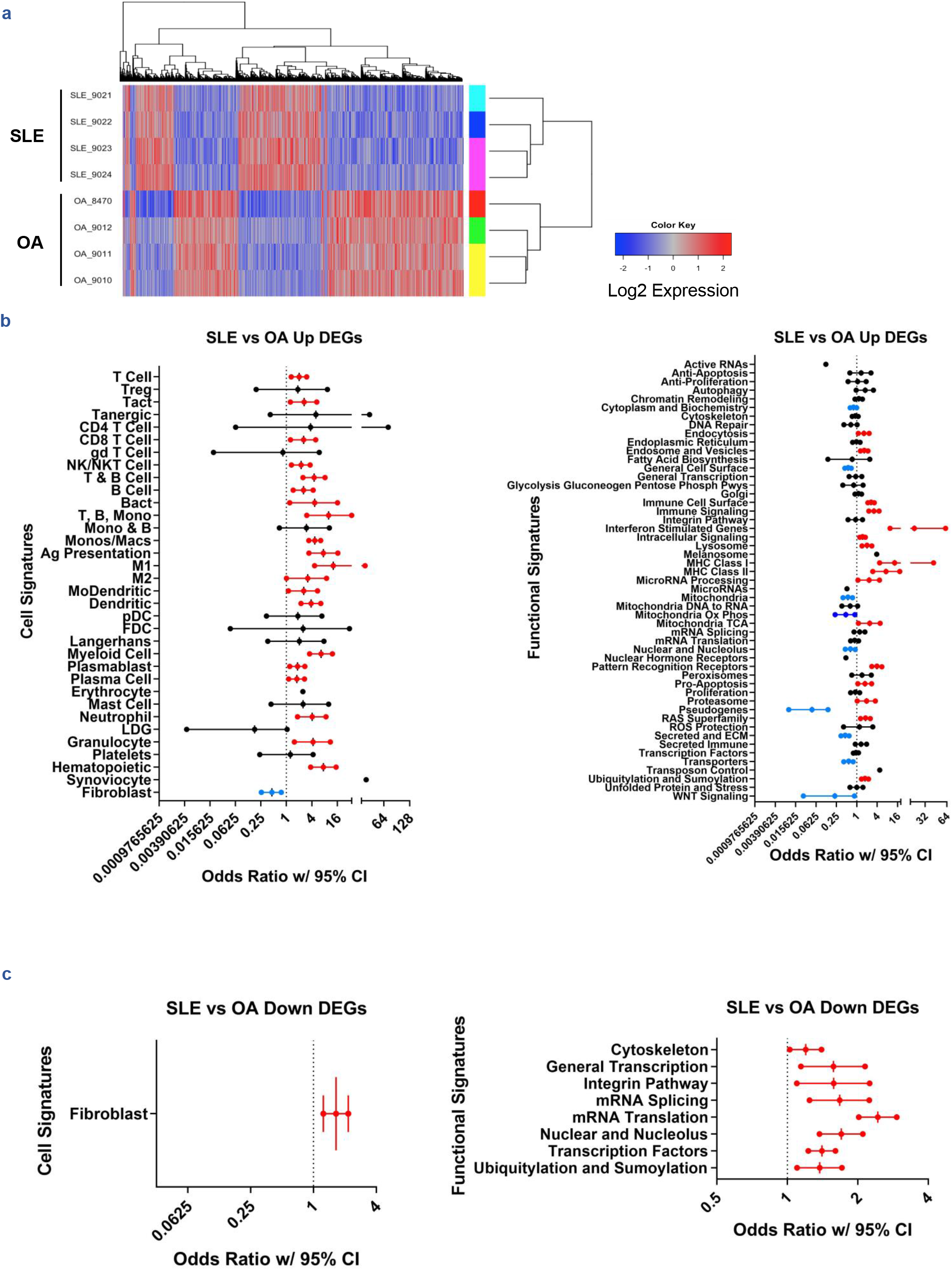
Overview of Gene Expression in SLE vs OA Synovium. **(a)** Heatmap of 6,496 DEGs from LIMMA analysis of SLE and OA synovial gene expression data. Increased **(b)** and decreased **(c)** transcripts were each characterized by cellular signatures for prevalence of specific cell types. DE transcripts were also characterized for functional signatures. Enrichment plots in **(b, c)** represent odds ratios bound by 95% confidence intervals (CI) using Fisher’s Exact Test. Significant enrichment by p-value (p<0.05) and confidence intervals that exclude odds ratio = 1 are colored red and blue for positive or negative association with the sample, respectively. The x-axes are plotted on log2 scales. For categories represented by a single point, odds ratio = 0 and the data point shown represents the upper bound of the confidence interval.

Of the 4,019 downregulated DEGs, only 17 were immune cell transcripts and thus downregulated genes did not reflect a change in hematopoietic cells (odds ratio of 0.0749, p=1). Notably, however, the fibroblast gene signature was downregulated (Fig. 1c). Functional analysis identified several molecular processes that were decreased in LA, most of which related to transcriptional activity/nuclear processes and cytoskeletal and integrin pathway changes. A list of genes significantly up- and downregulated in LA can be found in Supplementary Data S1 online.

Ingenuity Pathway Analysis (IPA) identified predominantly innate immune signaling processes, including a proinflammatory macrophage response, interferon signaling and inflammasome pathway activation, as well as an adaptive immune response (e.g., Th1 Pathway) (Fig. 2a). These were confirmed by functional analysis of IPA-predicted upstream regulators (UPRs) of the disordered gene expression profiles in LA (Fig. 2b), which identified specific intracellular signaling molecules, PRRs, and secreted immune proteins, including type 1 and 2 interferons. Of note, a number of pro-apoptotic genes, including *TNF, TNFSF10*, and *FAS* were predicted UPRs. The full export of IPA canonical pathway and upstream regulator data can be found in Supplementary Data S3-6 online.

**Figure 2.**
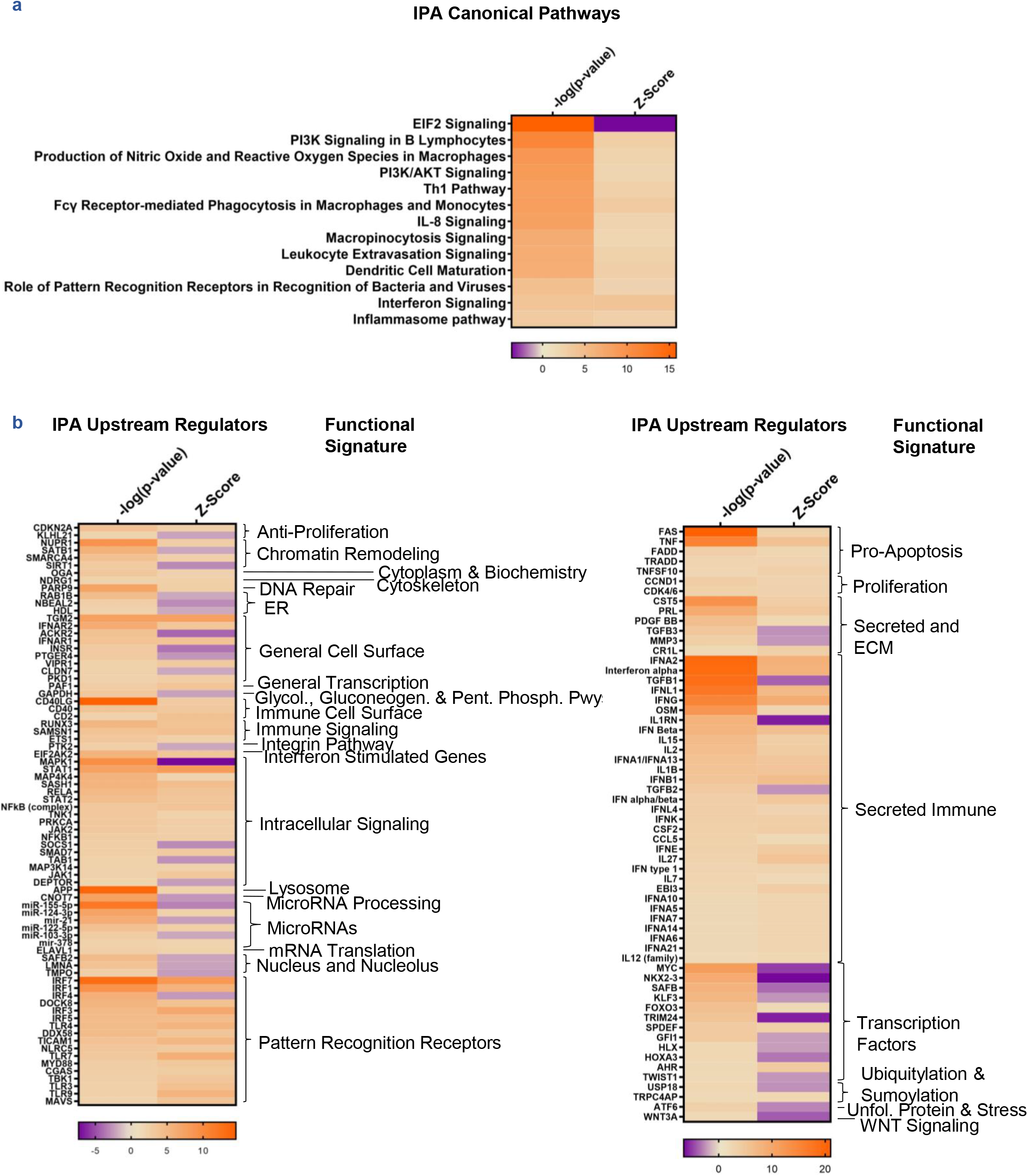
IPA-Predicted Signaling Pathways and Upstream Regulators Operative in LA. **(a)** Canonical pathways predicted by IPA based on DEGs, ordered by significance. **(b)** Significant upstream regulators predicted by IPA based on DEGs, ordered alphabetically by functional category. All canonical pathways and upstream regulators are significant by |Activation Z-Score| ≥ 2 and overlap p-value < 0.01.

To confirm DEG-identified molecular pathways and further interrogate LA gene expression, we implemented weighted gene co-expression network analysis (WGCNA), which serves as an orthogonal bioinformatic approach. WGCNA identified 52 modules, six of which were highly co-expressed and significantly associated with LA (see Supplementary Figs. S1, S2, and Supplementary Table S1 online). Of the six modules with significant, positive correlations to features of lupus, some correlated with the presence of LA and some modules correlated with SLEDAI and anti-dsDNA. The associations of the WGCNA modules with features of LA can be seen in Supplementary Figs. S1, S3, S4, and Supplementary Data S7-13 online. Many of the modules contained immune/inflammatory cell signatures. Notably, plasma cell genes were found in the midnightblue module and included IgG1, IgM, and IgD, indicating both pre- and postswitch plasmablasts/plasma cells as well as the presence of Igκ, Igλ, and numerous VL chains, signifying a polyclonal population (see Supplementary Fig. S5 online). Midnightblue, however, was upregulated in two of four lupus patients and negatively correlated with systemic disease (see Supplementary Fig. S1 online). The blue module, which was negatively correlated with LA, was enriched in synovial fibroblasts (see Supplementary Fig. S4, Supplementary Table S2 online), whereas the brown module, which was positively associated with LA, was less likely to be enriched for synovial fibroblasts (see Supplementary Fig. S3 online).

WGCNA modules were enriched in immune/inflammatory processes, including interferon stimulation, antigen presentation, immune cell surface markers, immune signaling, pathogenic pattern recognition, and lysosome function (see Supplementary Fig. S6 online), confirming BIG-C and IPA results. Further IPA canonical pathway analysis, IPA UPR analysis, and STRING/MCODE clustering of WGCNA modules (see Supplementary Fig. S7 online) indicated that the brown and navajowhite2 modules were characterized by myeloid cell responses, whereas the honeydew1 module was most characterized by interferon signaling and the darkgrey module with cellular activation, antigen presentation, and proinflammatory signaling. In contrast, the midnightblue module revealed T-cell: B-cell crosstalk, T-cell activation and differentiation, and B-cell signaling. Finally, salmon4 was not enriched in immune cells (see Supplementary Fig. S3 online).

### Lymphocyte Trafficking and GC Activity in LA

We assessed chemokine receptor-ligand pairs and adhesion molecules to understand mechanisms of immune/inflammatory cell localization in LA. Numerous chemokine receptorligand pairs were expressed in SLE synovium, including *CCR5-CCL4/5/8, CCR1-CCL5/7/8/23*, and *CXCR6-CXCL16* (Table 1). Of note, *CXCL13* was expressed in the midnightblue module and upregulated, although its receptor *CXCR5* was not detected in LA-associated WGCNA modules nor in DEGs. Adhesion molecules were found in LA-associated WGCNA modules including *VCAM1, CD44, CADM3*, and *ITGB2*.

**Table 1.** Chemokine Receptor-Ligand Pairs and Adhesion Molecules Associated with LA

| Gene Transcript | Name | SLE vs OA Analysis |  |
| --- | --- | --- | --- |
|  |  | DE LFC | WGCNA LA-associated module |
| CCL19 | Chemokine (C-C motif) ligand 19 | n/s | Honeydew1 |
| CCR2 | Chemokine (C-C motif) receptor 2 | 1.50 | Midnightblue |
| CCL2 | Chemokine (C-C motif) ligand 2 | n/s | Darkgrey |
| CCL7 | Chemokine (C-C motif) ligand 7 | n/s | Darkgrey |
| CCL8 | Chemokine (C-C motif) ligand 8 | 2.76 | Darkgrey |
| CCR5 | Chemokine (C-C motif) receptor 5 | 1.90 | Navajowhite2 |
| CCL4 | Chemokine (C-C motif) ligand 4 | 2.67 | Brown |
| CCL5 | Chemokine (C-C motif) ligand 5 | 1.94 | Midnightblue |
| CCL8 | Chemokine (C-C motif) ligand 8 | 2.76 | Darkgrey |
| CCR1 | Chemokine (C-C motif) receptor 1 | 2.14 | Brown |
| CCL5 | Chemokine (C-C motif) ligand 5 | 1.94 | Midnightblue |
| CCL7 | Chemokine (C-C motif) ligand 7 | n/s | Darkgrey |
| CCL8 | Chemokine (C-C motif) ligand 8 | 2.76 | Darkgrey |
| CCL23 | Chemokine (C-C motif) ligand 23 | 0.670 |  |
| CCR3 | Chemokine (C-C motif) receptor 3 | n/s | Darkgrey |
| CCL5 | Chemokine (C-C motif) ligand 5 | 1.94 | Midnightblue |
| CCL7 | Chemokine (C-C motif) ligand 7 | n/s | Darkgrey |
| CCL8 | Chemokine (C-C motif) ligand 8 | 2.76 | Darkgrey |
| CCRL2 | Chemokine (C-C motif) receptor-like 2 | 1.04 | Navajowhite2 |
| CKLF | Chemokine like factor | 0.297 |  |
| CMKLR1 | Chemokine-like receptor 1 | 1.59 | Darkgrey, Honeydew1 |
| CXCL2 | Chemokine (C-X-C motif) ligand 2 | 2.98 | Honeydew1 |
| CXCL3 | Chemokine (C-X-C motif) ligand 3 | 1.77 |  |
| CXCL8 | Chemokine (C-X-C motif) ligand 8 | 2.17 | Brown, Darkgrey |
| CXCR3 | Chemokine (C-X-C motif) receptor 3 | 1.45 | Midnightblue |
| CXCL9 | Chemokine (C-X-C motif) ligand 9 | 5.59 | Midnightblue |
| CXCL10 | Chemokine (C-X-C motif) ligand 10 | 4.81 | Midnightblue |
| CXCL11 | Chemokine (C-X-C motif) ligand 11 | 3.32 | Midnightblue |
| CXCR4 | Chemokine (C-X-C motif) receptor 4 | 1.39 | Brown |
| CXCL13 | Chemokine (C-X-C motif) ligand 13 | 3.47 | Midnightblue |
| CXCR6 | Chemokine (C-X-C motif) receptor 6 | n/s | Midnightblue |
| CXCL16 | Chemokine (C-X-C motif) ligand 16 | 0.768 | Navajowhite2 |
| CXCL11 | Chemokine (C-X-C motif) ligand 11 | 3.32 | Midnightblue |
| CX3CL1 | Chemokine (C-X3-C motif) ligand 1 | 0.453 |  |
| XCL1 | Chemokine (X-C motif) ligand 1 | n/s | Midnightblue |
| ALCAM | Activated leukocyte cell adhesion molecule | 1.55 |  |
| VCAM1 | Vascular cell adhesion molecule 1 | n/s | Navajowhite2 |
| CD44 | CD44 molecule | 1.25 | Brown, Darkgrey |
| ITGB1 | Integrin subunit beta 1 | -0.255 |  |
| ITGB2 | Integrin subunit beta 2 | 1.56 | Brown, Honeydew1 |
| ICAM1 | Intercellular adhesion molecule 1 | 0.861 | Darkgrey, Honeydew1, Midnightblue |
| ICAM3 | Intercellular adhesion molecule 3 | n/s | Midnightblue |
| PECAM1 | Platelet/endothelial cell adhesion molecule 1 | 0.618 | Salmon4 |
| SDK1 | Sidekick cell adhesion molecule 1 | -0.892 |  |
| SDK2 | Sidekick cell adhesion molecule 2 | -1.33 |  |
| CADM1 | Cell adhesion molecule 1 | -0.974 |  |
| CADM3 | Cell adhesion molecule 3 | n/s | Darkgrey |
| JAM2 | Junctional adhesion molecule 2 | -0.587 |  |
| JAM3 | Junctional adhesion molecule 3 | -0.673 |  |
| MCAM | Melanoma cell adhesion molecule | -1.08 |  |

DEGs and LA-associated WGCNA modules were assessed for adhesion molecules and
chemokine receptor-ligand pairs. Receptor-ligand pairs are grouped together in the table with
groupings alternately shaded. Log fold changes rounded to 3 significant figures are presented where available; otherwise, n/s=not significant.

We also examined expression of specific follicular helper T cell (T_fh_) and germinal center (GC) B-cell markers to determine whether GCs might contribute to LA pathogenesis (see Supplementary Fig. S8 online). *ICOS*, but not other T_fh_ markers, were found in LA. In addition, several GC B-cell markers were upregulated in SLE synovium, including, *CXCL13* and *IRF4*. However, *BCL6* and *RGS16* were notably downregulated and *RGS13* was not differentially expressed between SLE and OA. A cluster of GC B-cell markers that tended to be upregulated were co-expressed in the midnightblue module.

### GSVA Enrichment of Immune and Tissue Populations and Signaling Pathways

To assess the differences between SLE and OA synovitis on an individual sample basis, GSVA of various informative gene sets was carried out (Fig. 3, see Supplementary Data S14 online). Enrichment of hematopoietic cell types confirmed the presence of an immune infiltrate in LA, but not OA (Fig. 3a). Most cell types, including lymphoid and myeloid populations, were enriched. Cytokine signaling was also enriched in LA, including both proinflammatory cytokine signaling as well as inhibitory cytokines (Fig. 3b). Of note, the downstream signature induced by TNF signaling was significantly enriched in LA (p=0.00918). Whereas antigen presentation markers, cellular activation markers, and the inflammasome pathway were enriched in LA compared to OA, a cell cycle/proliferation signature and complement pathways were not enriched (Fig. 3c). Along with upregulation of inhibitory cytokines, inhibitory receptors and negative regulation of T cells were enriched in LA (Fig. 3d). Most of the previously noted IPA-predicted canonical signaling pathways were enriched in SLE synovium aside from signaling by the eukaryotic initiation factor eIF2 (Fig. 3e).

**Figure 3.**
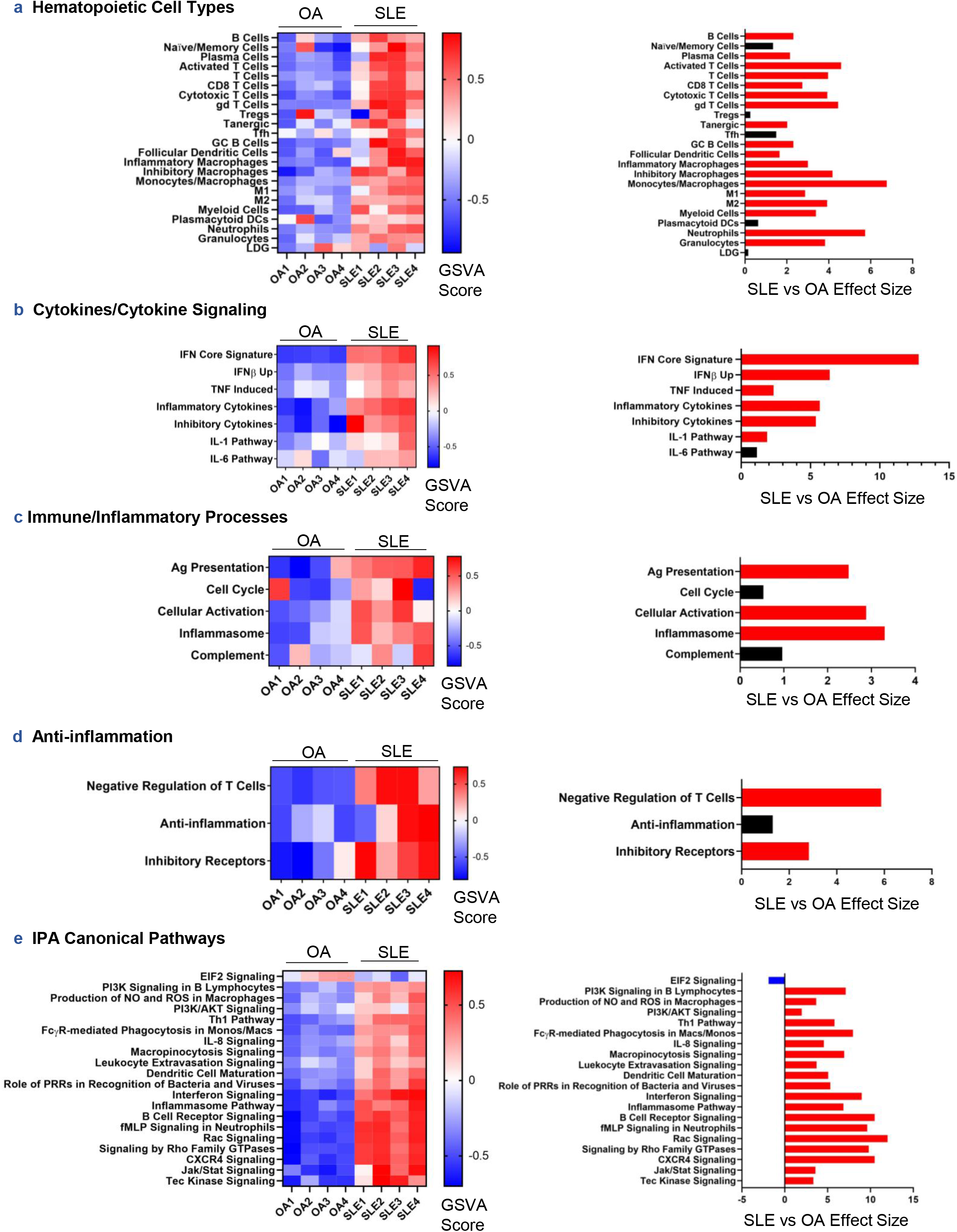
GSVA Enrichment of Immune/Inflammatory-Specific Cells and Processes in LA. GSVA of hematopoietic cell types **(a)**, cytokine signatures and signaling pathways **(b)**, immune/inflammatory processes **(c)**, anti-inflammatory processes **(d)**, and IPA-predicted canonical signaling pathways from DEGs in SLE vs OA synovium **(e)** was conducted on log2-normalized gene expression values from OA and SLE synovium. Hedge’s g effect sizes were calculated with correction for small sample size for each gene set and significant differences in enrichment between cohorts were found by Welch’s t-test (p<0.05), shown in the panels on the right. Red and blue effect size bars represent significant enrichment in SLE and OA, respectively.

Conversely, OA synovium was enriched in tissue repair/destruction and markers of fibroblasts (Fig. 4a). Querying LA and OA synovial expression profiles with the co-expressed genes from fibroblast subpopulations described in human RA and OA synovium [10], we found that two resident synovial sublining fibroblast populations, CD34^+^ and DKK3^+^, were significantly increased in OA over LA (Fig. 4b; p=0.00148 and 0.00213, respectively). However, co-expressed genes characterizing the HLA-DR^hi^ sublining fibroblast population were significantly enriched in LA (p=2.70e-05). Interestingly, using the same approach to assess macrophage populations also described in human RA and OA [10–11], we found quiescent macrophages and IFN-activated macrophages significantly increased in LA (Fig. 4c; p=0.0332 and 1.14e-06, respectively), whereas phagocytic macrophages were associated with OA. HBEGF^+^ proinflammatory macrophages tended to be enriched in LA but did not reach statistical significance (p=0.0993).

**Figure 4.**
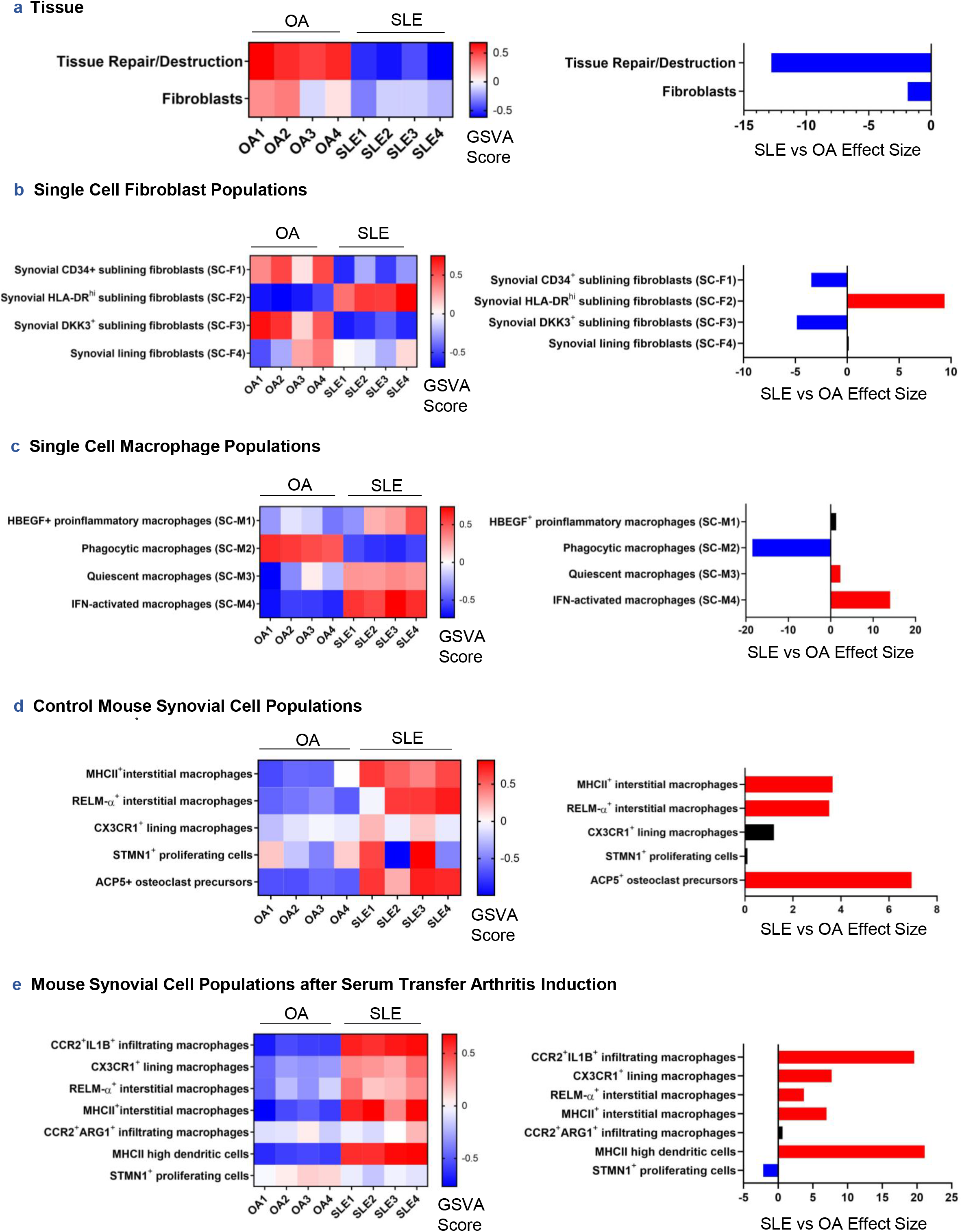
GSVA Enrichment of Tissue-Specific Cells and Processes in LA. GSVA of synovial tissue processes and specific cell types **(a)** and recently published synoviumspecific cell subtypes in human RA, OA, and mouse synovium **(b-e)** was conducted on log2-normalized gene expression values from OA and SLE synovium. Hedge’s g effect sizes were calculated with correction for small sample size for each gene set and significant differences in enrichment between cohorts were found by Welch’s t-test (p<0.05), shown in the panels on the right. Red and blue effect size bars represent significant enrichment in SLE or OA, respectively. Literature-derived signatures in **(b-e)** underwent co-expression analyses before being used as GSVA gene sets (see Methods).

Other unique macrophage subpopulations have been described in mouse synovium, including a CX3CR1^+^ resident subtype resembling epithelial cells [12]. Taking the co-expressed genes of the human orthologs of this macrophage signature and others detected in healthy mouse synovium, we found that two types of interstitial macrophages, including MHCII^+^ and RELM-α^+^ populations, were significantly more abundant in LA (p=0.00549 and 0.00802, respectively) as well as a group of ACP5^+^ osteoclast precursors (Fig. 4d; p=9.37e-04). The CX3CR1^+^ lining macrophage population from healthy mice was not significantly enriched in either LA or OA synovium (p=0.110); however, the CX3CR1^+^ lining macrophage population from murine inflammatory arthritis were increased in LA (Fig. 4e). Additionally, inflammatory arthritis-associated MHCII^+^ and RELM-α^+^ interstitial macrophages were enriched in LA as were MHCII^hi^ dendritic cells and CCR2^+^IL1B^+^ monocyte-derived macrophages.

### Comparison of Gene Expression in LA and RA Synovitis

Fewer genes were upregulated in RA, but 18% of these genes identified immune/inflammatory cells compared with 10% of genes upregulated in LA (Fig. 5a). Characterization of upregulated DEGs by cell signatures revealed greater numbers of myeloid and monocyte/macrophage-specific transcripts in LA compared to RA, whereas immune infiltrates in RA were more characteristic of T- and B-cells. B-cells, naïve/memory cells, and gamma delta (gd) T cells were significantly increased in RA over LA (p=0.0249, p=0.0121, and p=0.00689, respectively), whereas monocytes/macrophages, inhibitory macrophages, and M2 macrophages were significantly increased in LA over RA (Fig. 5b; p=0.0160, p=0.00306, and p=0.00719, respectively).

**Figure 5.**
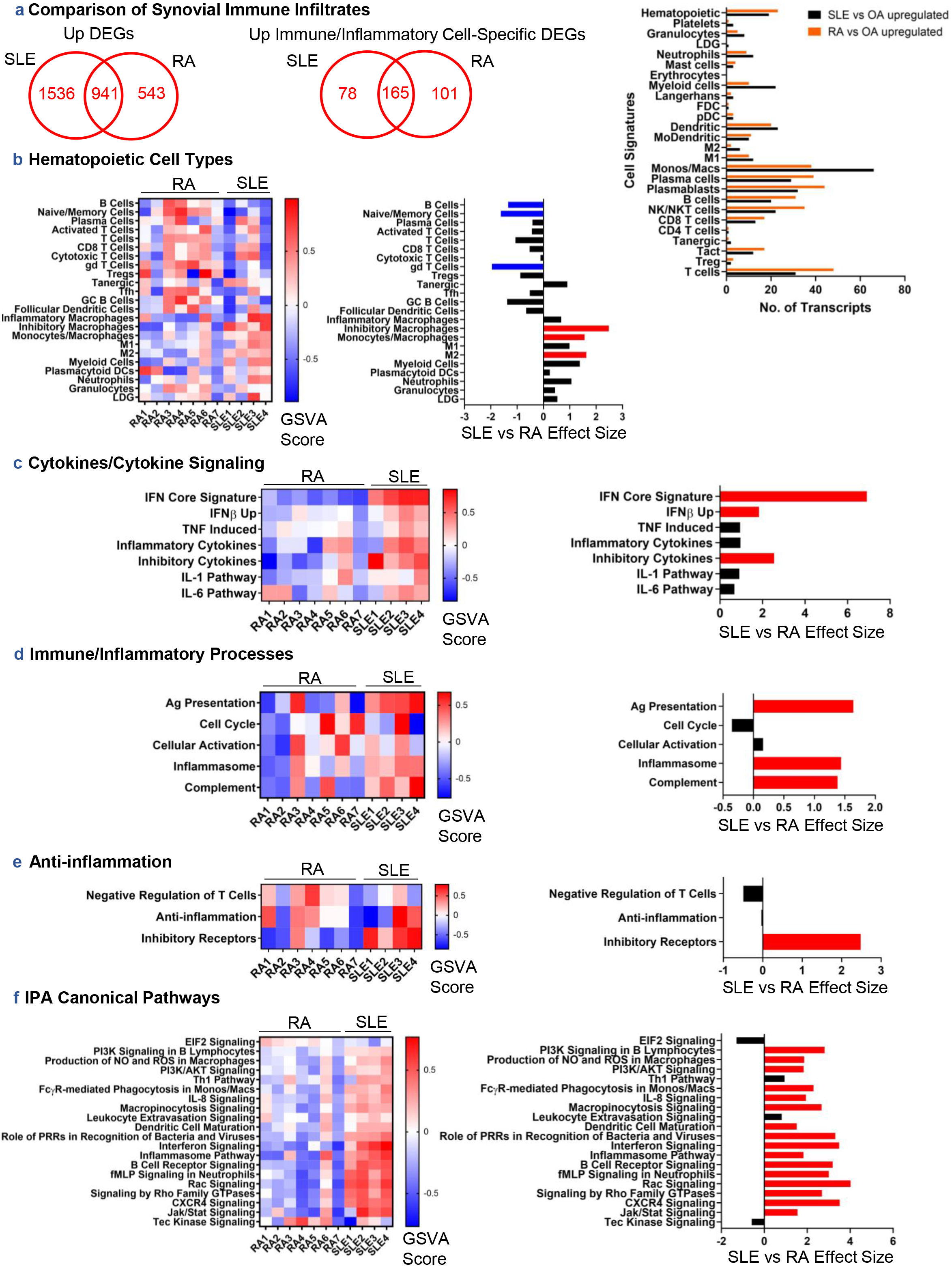
Comparison of Gene Expression in SLE and RA Synovitis. A comparison of immune/inflammatory gene signatures between SLE and RA synovium using 7 RA patients from GSE36700. **(a)** Upregulated DEGs were identified between RA and OA synovium, compared to DEGs from SLE vs OA synovium, and characterized by cellular signatures. GSVA of hematopoietic cell types **(b)**, cytokine signatures and signaling pathways **(c)**, immune/inflammatory processes **(d)**, anti-inflammatory processes **(e)**, and IPA-predicted canonical signaling pathways from DEGs in SLE vs OA synovium **(f)** was conducted on log2-normalized gene expression values from SLE and RA synovium. Hedge’s g effect sizes were calculated with correction for small sample size for each gene set and significant differences in enrichment between cohorts were found by Welch’s t-test (p<0.05), shown in the panels on the right. Red and blue effect size bars represent significant enrichment in SLE or RA, respectively.

Interferon signaling pathways, the complement pathway, inflammasome pathway, antigen presentation, inhibitory cytokines, and inhibitory receptors were substantially enriched in LA, whereas negative regulation of T-cells and cell cycle tended to be enriched in RA (Fig. 5c-e). Downstream signatures of TNF, IL-1, and IL-6 tended to be enriched in LA compared to RA. Finally, other pathways, including response to stimuli, phagocytosis, chemokine signaling, BCR signaling, and PI3K signaling, were considerably enriched in LA (Fig. 5f).

Transcripts associated with tissue repair/destruction were significantly enriched in RA (see Supplementary Fig. S9 online; p=9.95e-04). Enrichment of general fibroblast markers was not uniform in either tissue. Nonetheless, HLA-DR^hi^ sublining and CD55^+^ lining fibroblasts were significantly associated with LA (p=0.00247 and p=1.39e-04, respectively), whereas the CD34^+^ and DKK3^+^ sublining populations were depleted in LA compared to RA.

HBEGF^+^ proinflammatory macrophages and phagocytic macrophages tended to be more enriched in RA patients, although not uniformly, whereas quiescent and IFN-activated macrophages were more enriched in LA, the latter population reaching statistical significance (see Supplementary Fig. S9 online; p=0.00857) [10–11]. When detected, the normal and arthritis murine equivalent macrophage populations trended towards enrichment in LA, including the CX3CR1^+^ lining macrophages [12].

### Compounds Predicted to Target LA

Drugs predicted to reverse the abnormal gene expression profile of LA were identified by connectivity mapping to the LINCS database and are shown in Table 2 and Supplementary Table S3 online. Most abundantly predicted compounds include anti-cancer drugs targeting tubulin polymerization, MAPK signaling, and EGFR signaling, as well as current lupus standard-of-care therapies, including corticosteroids and NSAIDs/prostaglandin synthesis inhibitors. Interestingly, a few alternative medicines were predicted to counteract LA, including curcumin, capsaicin, resveratrol, and caffeine. In addition to the LINCS-predicted compounds, we sought to expand the potential list of therapies to include those targeting biological upstream regulators (BURs). The top 50 BURs determined by connectivity scoring with gene expression generated by knock down or overexpression studies in cell lines are summarized in Fig. 6a along with drugs that could potentially directly target these BURs, including ESR1, type 1 IFN, bromodomain and casein kinase inhibitors. Finally, the UPRs predicted by IPA were also matched with potential targeting drugs (Fig. 6b). Notably, 26% of drugs targeting IPA upstream regulators were also predicted by LINCS BURs drug-target matches (see Supplementary Data S15-16 online), and included inhibitors of TNF, type I interferon, the NFκB pathway, JAK, and CDK.

**Figure 6.**
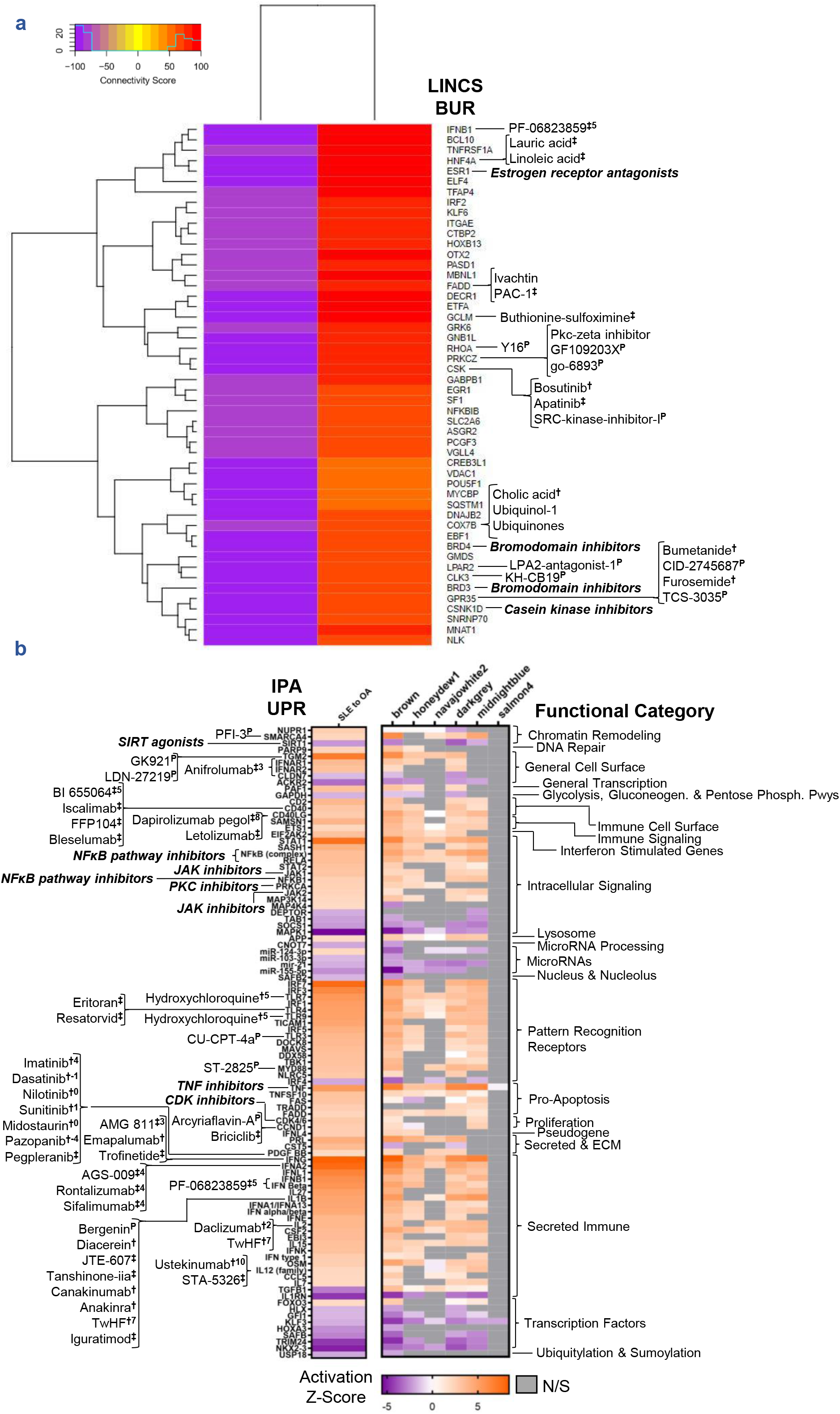
LINCS Biological Upstream Regulators and IPA Upstream Regulators Operative in LA Are Potential Druggable Targets. **(a)** The top 50 targets (BURs) opposing the LA gene signature from LINCS knock down (KD) and overexpression (OE) assays summarized by connectivity score and matched to appropriate targeting drugs. KD and OE data were filtered for connectivity scores in the [−75 to −100] and [50 to 100] ranges, respectively. **(b)** The consensus IPA-predicted UPRs between DEGs and LA-associated WGCNA modules summarized by Activation Z-Score, functional category, and also matched to appropriate targeting drugs. Drugs and compounds targeting the BURs and UPRs were sourced from LINCS/CLUE, IPA, literature mining, CoLTS [41], STITCH, and clinical trials databases. Drug annotations are grouped together by target and CoLTS scores (range −16 to +11) are displayed as integers in superscript. Some upstream regulators are matched to groups of drugs (e.g., NFκB pathway inhibitors, bold, italicized), for which the full list of drug-target matches can be found in Supplementary Data S15-16 online. ^P^: Preclinical; ^‡^: Drug in development/clinical trials; ^†^: FDA-approved

**Table 2.**
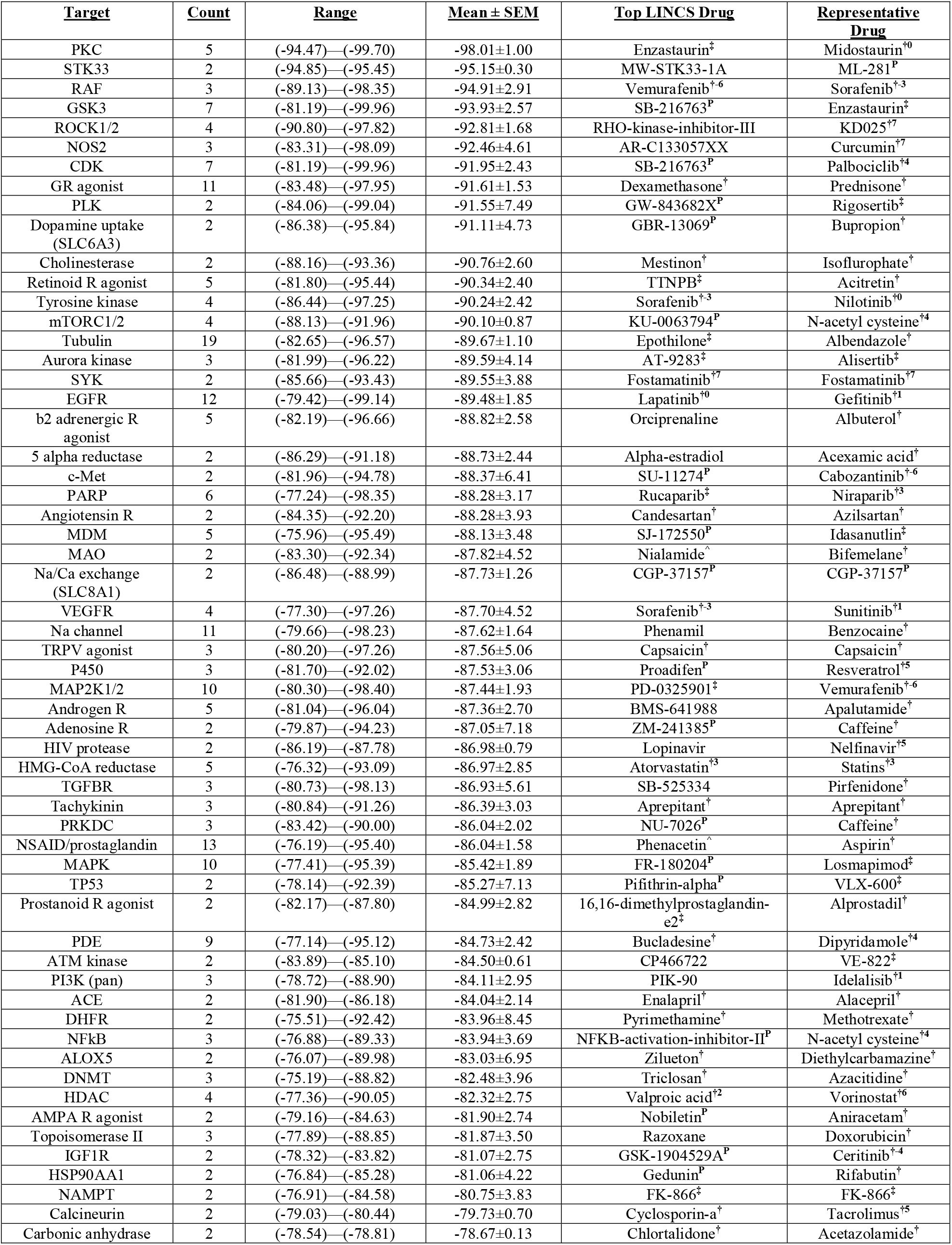
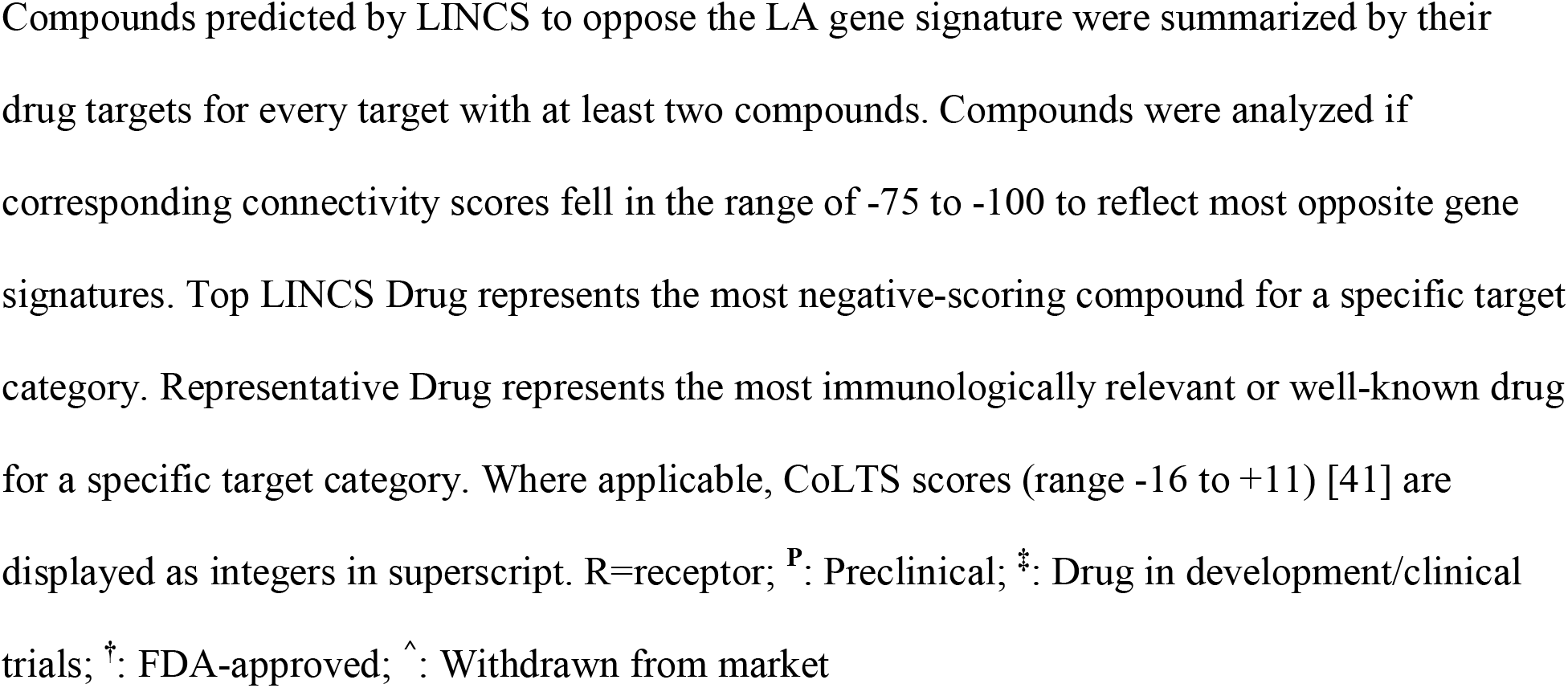
Compounds Targeting LA

## DISCUSSION

Analyses of DEGs revealed a uniform inflammatory infiltrate in LA of mostly myeloid lineage cell types, including monocytes, M1 macrophages, antigen presenting cells, and other myeloid and hematopoietic cells. WGCNA indicated enrichment of other immune cell types, including activated and effector T cells, NK cells, B cells, plasma cells/plasmablasts, and both M1 and M2-polarized macrophages, suggesting both innate and adaptive mechanisms at play in LA, although the involvement of adaptive immune cells was less uniform than that of the innate immune system.

Our findings indicated that myeloid-lineage cells were enriched in LA and, therefore, may play a central role in the observed inflammation. IPA revealed monocyte/macrophage-mediated phagocytosis and NO and ROS production signaling pathways, and GSVA confirmed gene expression profiles of both inflammatory M1 and inhibitory M2 macrophages in LA. This aligns with prior histology suggesting the presence of infiltrating macrophages [9] and recent analysis of myeloid cells in SLE blood associated an M1 inflammatory phenotype with active versus inactive disease [13]. By comparison to single-cell transcriptional profiles from sorted murine synovial CD45^+^Cd11b^+^Ly6G^-^ cells [12], significant enrichment of resident macrophage populations as well as non-resident infiltrating populations were identified, as well as antiinflammatory macrophage subpopulations and inhibitory and inflammatory cytokines, indicating that multiple macrophage subtypes and their secreted products may contribute to and perhaps protect against LA pathogenesis.

Fibroblast-unique genes were downregulated in LA, possibly representing local relative loss or diminished/altered function of resident fibroblasts. Pathologic fibroblast populations, potentially contributing to local tissue damage, have been shown to reside in the synovium of patients with leukocyte-rich RA [14–16], including a subpopulation of CD34^+^ sublining fibroblasts, which was decreased in LA. Another fibroblast population enriched in leukocyte-rich RA and characterized by higher expression of MHC Class II genes *IL6* and *CXCL12* appeared to also be enriched in LA [10]. However, this population was mainly characterized by interferon stimulated and MHC Class I/II genes, and, therefore, this signature cannot definitively be attributed to fibroblasts (see Supplementary Fig. S10 online). Furthermore, comparison of SLE and RA synovial gene profiles indicated maintenance of a lining CD55^+^ fibroblast layer in LA and tissue repair/destruction mechanisms in RA. LA may, therefore, differ from RA and OA, in which joint organ pathology is characterized by fibroblast-mediated tissue damage and, rather, be characterized by a loss of function or dysregulation of proinflammatory fibroblasts with destructive potential.

LA may also differentiate from RA in its immune cellularity and composition. A greater number of genes were found significantly altered in LA than in RA but a smaller portion of these transcripts could be attributed to immune/inflammatory cell populations, indicating an overall greater immune infiltrate in RA than in LA. Of the immune/inflammatory cell-specific transcripts identified, RA upregulated DEGs indicated increased T-cells, B-cells, NK/NKT-cells, and other lymphocytes, whereas LA upregulated DEGs were more characteristic of monocytes/macrophages and myeloid cells. Thus, LA may be more myeloid-mediated than RA. GSVA replicated this finding with significant upregulation of the core type I interferon signature, antigen presentation signature, inflammasome pathways, and monocyte/macrophage cell populations in LA including, notably, more inhibitors of inflammation. Specific macrophage and dendritic cell subsets originally identified in mouse synovium [12] were also more enriched in LA. Interestingly, the downstream TNF, IL-1, and IL-6 signatures tended to be more enriched in LA than RA, indicating potential for repurposing anti-TNF biologics, the IL-1 antagonists anakinra and canakinumab, and IL-6R antagonist tocilizumab to treat LA.

The detection of Ig heavy chain pre- and post-switch plasma cells in LA was notable. *IRF4, XBP1*, and *PRDM1*, genes essential for plasma cell maturation [17], were all detected in LA. There was some evidence of GC formation in LA, but *BCL6, AICDA*, and *RGS13* were not upregulated nor detected in an LA-associated WGCNA module. However, *CXCL13*, encoding a chemoattractant that has been reported in RA synovial GCs [18], was strongly upregulated. These findings suggest that fully-developed GCs are not a routine part of LA. Rather, it is more likely that lupus synovium contains lymphoid aggregates that support B-cell proliferation and autoantibody formation, as reported in the spleen in immune thrombocytopenia [19]. An interesting caveat to our data is the strong negative correlations of the midnightblue module, which contains the plasma cell signature, to SLEDAI and anti-dsDNA, whilst being positively correlated to LA. This suggests that the presence of plasmablasts/plasma cells in lupus synovium may not contribute significantly to systemic autoantibody levels and extra-articular lupus disease activity. Rather, the nature of the local inflammation may facilitate entry of circulating plasmablasts/plasma cells into the synovial space and/or their local differentiation.

The overexpression of numerous chemokines and chemokine receptors suggests chemokine signaling may play an important role in the infiltration of immune/inflammatory cells into lupus synovium. *CXCR3* and its ligands *CXCL9, CXCL10*, and *CXCL11* were all found upregulated and co-expressed in the midnightblue module, which contained a robust lymphocyte signature. This signaling axis is known to be induced by IFNγ and is involved in the recruitment of activated lymphocytes, particularly of naïve T cells and their differentiation into Th1 cells [20]. *CXCR3* and *CXCR4*, both of which were upregulated in LA, are additionally important for the homing and maintenance of plasma cells [21]. These chemokine receptors could be involved in the recruitment of circulating plasmablasts/plasma cells into lupus synovium and/or their *in situ* retention and/or differentiation [22–24]. Other chemokines and their receptors such as *CCR5-CCL4/CCL5* could contribute to recruitment of other leukocytes into the synovium, including macrophages, monocytes, and T-cells [25].

We utilized gene expression analysis to predict novel drugs that might target abnormally expressed genes or pathways and suppress inflammation. Predicted drugs and compounds identified novel potential therapies, but also confirmed current treatments by identifying standard-of-care lupus drugs such as glucocorticoids, methotrexate, aspirin, and cyclosporine. Notably, a large number of anti-cancer drugs with variable mechanisms of action were also predicted.

Drugs targeting the CDK family were predicted to revert the LA gene signature and may point to potential repurposing of drugs such as palbociclib or related seliciclib and other CDK inhibitors that have been shown to ameliorate nephritis in animal models [26], possibly by reducing proliferation of lupus T- and B-cells *in vitro* [27]. Similarly, bucladesine was one of nine phosphodiesterase inhibitors predicted to suppress LA. Other novel druggable therapeutic targets include ROCK2, SYK, PARP1 and PARP2, and HDAC.

Notably, a large number of sodium channel blockers were predicted to target LA, possibly related to increased nervous innervation of the inflamed synovium. Neurologic targets included the acetylcholine, dopamine, serotonin, GABA-A, adrenergic, and glutamate receptors. These may have been predicted based on changes in the innervation of the inflamed tissue, although an effect on immune/inflammatory cells is also possible [28]. Similarly, transmembrane ion channels were predicted targets of the LA gene program and may reflect dysregulation of innervation or a role on immune/inflammatory cells.

Multiple drug predictions were made surrounding the estrogen and progesterone receptors. It is well-known that women are affected by systemic lupus 10 times more often than men and that sex hormones are involved in modulating the immune system [29]. While glucocorticoids have mainly anti-inflammatory and immunosuppressive effects, sex hormones such as estrogen and progesterone may have either pro-inflammatory or anti-inflammatory effects depending on the types of receptors expressed and other factors [30]. Estrogen may increase risk of disease by favoring autoreactive B-cells and promoting type I interferon production, whereas progesterone seems to counteract these effects [31]. Thus, the right balance of these hormones may attenuate disease activity. In an all-female cohort, we found *ESR1*, encoding the alpha estrogen receptor, to be an upstream regulator of LA. Tamoxifen was also repeatedly suggested as a potential therapy for LA and has shown utility in murine lupus and in human lupus T-cells [32–34]. However, no clinical trials of Tamoxifen have been conducted in lupus or other autoimmune disease and cases of Tamoxifen-induced lupus and other adverse outcomes have been reported [35–36]. Thus, it is clear that female hormone receptors are important in LA pathogenesis but delineating their specific roles and the crosstalk between glucocorticoids and sex hormones needs further study.

Bioinformatic analysis of LA revealed a pattern of immunopathogenesis in which myeloid cell-mediated inflammation dominates. The breadth of the immune response underlying LA provides a basis for multiple avenues of therapeutic intervention to be considered that mouse models and previous studies have not provided. With these findings we can begin to hypothesize specific candidate target genes and pathways from which to develop or repurpose drugs to treat and improve LA specifically.

## METHODS

### Gene Expression Data Sourcing and Patient Characteristics

Publicly available gene expression data from synovial biopsies from the knees of 4 SLE, 5 OA, and 7 RA subjects with active articular disease were obtained from NCBI Gene Expression Omnibus (GEO) under accession GSE36700 [8]. The SLE patients assessed had a mean (±s.d.) SLEDAI of 8.25 (1.71), CRP of 12.5 mg/L (4.12), C3 of 82.5 mg/dL (28.0), C4 of 13 mg/dL (3.56) and anti-dsDNA of 97.6 IU/mL (77.0), and all patients had active arthritis at the time biopsy was taken. Complete patient data can be found in Supplementary Data S17 online. Data processing and analysis were conducted within the R statistical programming platform using relevant Bioconductor packages.

### Data Normalization

All raw data files underwent background correction and GCRMA normalization resulting in log2 intensity values compiled into expression set objects (e-sets). Outliers were identified through the inspection of first, second, and third principal components and through inspection of array dendrograms calculated using Euclidean distances and clustered using average/UPGMA agglomeration. GSM899013_OA5 was consistently identified as an outlier and excluded from further analyses. Low intensity probes were removed by visual assignment of a 2.34 threshold cutoff upon a histogram of binned log2-transformed probe intensity values.

### Differential Gene Expression

Identification of DEGs in SLE vs OA samples (n=8) and RA vs OA samples (n=11) was conducted using the LIMMA package in R. To increase the probability of finding DEGs, both Affy chip definition files (CDFs) and BrainArray CDFs were used to created and annotate e-sets, analyzed separately, then results merged. Linear models of normalized gene expression values were created through empirical Bayesian fitting. Resultant p-values were adjusted for multiple hypothesis testing using the Benjamini-Hochberg correction. Significant probes were filtered to retain a pre-specified False Discovery Rate (FDR) < 0.2 and duplicate probes were removed again to retain the most significant probe. The FDR was assigned a priori to avoid excluding false negative probes. The full list of DEGs can be found in Supplementary Data S1-2 online.

### Weighted Gene Co-expression Network Analysis (WGCNA)

The same normalized and filtered data (Affy CDFs only) were inputted into WGCNA to conduct an unsupervised clustering analysis yielding statistically co-expressed modules of genes used for further biological interrogation. A scale-free topology matrix (TOM) was calculated to encode the network strength between probes. TOM distances were used to cluster probes into WGCNA modules. Resulting co-expression networks were trimmed using dynamic tree cutting and the deepSplit function in R. Partitioning around medoids (PAM) was also utilized to assign outliers to the nearest cluster. Modules were given random color assignments and expression profiles summarized by a module eigengene (ME). Final membership of probes representing the same gene were decided based on strongest within-module correlation to the ME value. For each module, ME values were correlated by Pearson correlation to clinical data including cohort, SLEDAI, anti-dsDNA, C3, C4, and CRP levels. Cohort was represented as a binary variable where SLE=1 and OA=0 whereas the remaining clinical data were continuous variables. Full module gene lists can be found in Supplementary Data S8-13 online.

### Quality Control (QC) and Selection of WGCNA Modules

WGCNA modules of interest underwent a QC process to ensure modules were reflective of disease state. First, ME expression per patient was visually inspected to assess consistency of gene expression in a given cohort. Second, module membership, or eigengene-based connectivity (k_ME_), was plotted against probe correlation to the primary clinical trait of interest (SLEDAI) to gauge how well a given module agreed to the clinical trait. Finally, the Pearson correlations of MEs to the clinical metadata were examined. Absolute values of correlation coefficients in the range 0.5-1 were considered strong and alpha=0.05 determined significance.

### Functional Analysis

Immune/Inflammation-Scope (I-Scope) and Biologically Informed Gene Clustering (BIG-C) are functional aggregation tools for characterizing immune cells by type and biologically classifying large groupings of genes, respectively. I-Scope categorizes gene transcripts into a possible 32 hematopoietic cell categories based on matching 926 transcripts known to mark various types of immune/inflammatory cells. BIG-C sorts genes into 52 different groups based on their most probable biological function and/or cellular/subcellular localization. Tissue-Scope (T-Scope) is an additional aggregation tool to characterize cell types found in specific tissues. In these analyses only the two T-Scope categories relevant to the synovium were used: fibroblasts and synoviocytes.

### Network Analysis

Cytoscape (V3.6.1) software was used to visualize protein-protein interactions based on the Search Tool for the Retrieval of Interacting Genes/Proteins (STRING) database via the stringApp plugin application. A confidence score of 0.40 was used. The clustermaker2 plugin application was used to created MCODE clusters of interrelated genes using a network scoring degree cutoff of 2, node score cutoff of 0.2, maximum depth of 100, and k-Core of 2. Genes not recognized by the STRING database were removed from datasets prior to upload into Cytoscape.

### Ingenuity ^®^ Pathway Analysis (IPA)

The canonical pathway and UPR functions of IPA core expression analysis (Qiagen) were used to interrogate DEGs and WGCNA module gene lists. Core expression analyses were based on fold change if uploaded genes were differentially expressed; otherwise, a fold change of one was used. Canonical pathways and UPRs were considered significant if Activation Z-Score| ≥ 2 and overlap p-value ≤ 0.01.

### Gene Set Variation Analysis (GSVA)

The GSVA R package was used as a non-parametric, unsupervised gene set enrichment method. Enrichment scores were calculated using a Kolgomorov Smirnoff (KS)-like random walk statistic to estimate variation of pre-defined gene sets. The inputs for the GSVA algorithm were e-sets containing log2 microarray expression values (Affy HGU133plus2 definitions) and predefined gene sets co-expressed in SLE datasets. Low-intensity probes were filtered out based on interquartile range (IQR) [37]. GSVA was conducted on the remaining network and Welch’s t-test was used to detect significant differences in enrichment between cohorts at an alpha level of 0.05, followed by calculation of Hedge’s g effect size with correction for small samples. Welch’s t-test was used to account for unequal variances in both the SLE and OA populations and RA and OA populations.

Enrichment gene sets containing cell type- and process-specific genes listed in Supplementary Data S14 online were created through an iterative process of identifying DE transcripts pertaining to a restricted profile of hematopoietic cells in 13 SLE microarray datasets and checked for expression in purified T cells, B cells, and monocytes to remove transcripts indicative of multiple cell types. Genes were identified through literature mining, GO biological pathways, and STRING interactome analysis as belonging to specific categories [38]. Select gene sets (i.e., TNF-induced, IFNβ-upregulated, M1 and M2) were derived directly from in vitro experiments [39–40]. The M1 signature was edited to remove interferon stimulated genes. Additionally, IL-1 and IL-6 gene sets were derived from the first three tiers of the respective PathCards signaling pathways.

### Co-expression Analysis

Co-expression analyses of literature-derived signatures published in mouse and human synovium were conducted in R. Briefly, Spearman’s rank correlation coefficients and p-values were computed using the rcorr() function based upon input log2 expression values for each gene in each SLE and OA sample. Spearman correlations were chosen to avoid assuming linear relationships. The input gene signature was refined to contain genes significantly correlating with at least 25% of the original gene signature at an alpha level of 0.05, then refined again to contain genes positively correlated with at least 25% of the new signature (i.e., Spearman’s rho>0). The final co-expressed signatures were used as GSVA gene sets. Mouse to human ortholog conversion was done using the homologene R package.

### LINCS Drug-Target Prediction and Biological Upstream Regulator Analysis

The Library of Integrated Network-Based Cellular Signatures (LINCS) perturbation database (http://data.lincscloud.org.s3.amazonaws.com/index.html) was queried using a DEG list of significantly up- and down-regulated genes from the SLE and OA samples. Comparisons were made based on LINCS-computed connectivity scores, where −100 describes a transcriptional program perfectly opposing the user-uploaded gene signature and 100 describes a transcriptional program perfectly representative of the user-uploaded gene signature.

### Drug-Target Matching

In addition to the LINCS-predicted compounds, LINCS-predicted biological upstream regulators (BURs) and IPA-predicted UPRs were annotated with respective targeting drugs and compounds to elucidate potential useful therapies in lupus synovitis. Drugs targeting gene products of interest both directly and indirectly were sourced by IPA, the Connectivity Map via the drug repurposing tool, GeneCards, STITCH (V5.0), Combined Lupus Treatment Scoring (CoLTS)-scored drugs [41], FDA labels, DrugBank, literature mining, and queries of clinical trials databases. The CLUE drug repurposing tool was accessed at https://clue.io/repurposing-app.

### STITCH

The Search Tool for Interactions of Chemicals (STITCH) (V5.0) database (http://stitch.embl.de/) of known and predicted protein-protein and protein-chemical interactions was used to predict direct and indirect drug targeting mechanisms. For each gene product of interest, the top 10 interactors were analyzed and drugs directly targeting the top interactors were matched according to the methods described. A medium confidence score cutoff of 0.4 for interaction predictions was used. Predicted interactions based solely on text-mining were not considered.

### Statistical Analysis

Enrichment statistics in SLE vs OA were calculated by right-sided Fisher’s Exact Test in R using the function fisher.test(). Statistical significance was obtained at p<0.05.

## DATA AVAILABILITY

The dataset analyzed during the current study is available in the NCBI GEO repository, https://www.ncbi.nlm.nih.gov/geo/query/acc.cgi?acc=GSE36700. Additional data generated from analyses are included in this published article (and its Supplementary Information files).

## ACKNOWLEDGEMENTS

The authors would like to thank Drs. FA Houssiau and BR Lauwerys who provided clinical data on the LA subjects studied in this report, Dr. Adam C. Labonte for his consultation on macrophage polarization and Dr. Kathryn Kingsmore Allison for her assistance with preparation of the manuscript and consultation on proper statistical technique.

The authors would additionally like to acknowledge prior presentation of this work as posters at the ACR Annual Meetings 2018 and 2019 [42–43].

## AUTHOR CONTRIBUTIONS

E. L. H. contributed to the planning and data analysis of the work described and wrote the main manuscript. M. D. C. and S. H. contributed to the planning and data analysis. P. B. and N. S. G. contributed to data analysis and software. R. R. contributed software. A. C. G. and P. E. L. contributed to the concept and design of the study, interpretation of the data, and revision and editing of the manuscript.

