## Supplementary Figures and Tables for "Analysis of Gene Expression from Systemic Lupus Erythematosus Synovium Reveals a Profile of Activated Immune Cells and Inflammatory Pathways"

### Supplementary Figure S1. WGCNA Reveals Six LA-Associated Modules of Genes

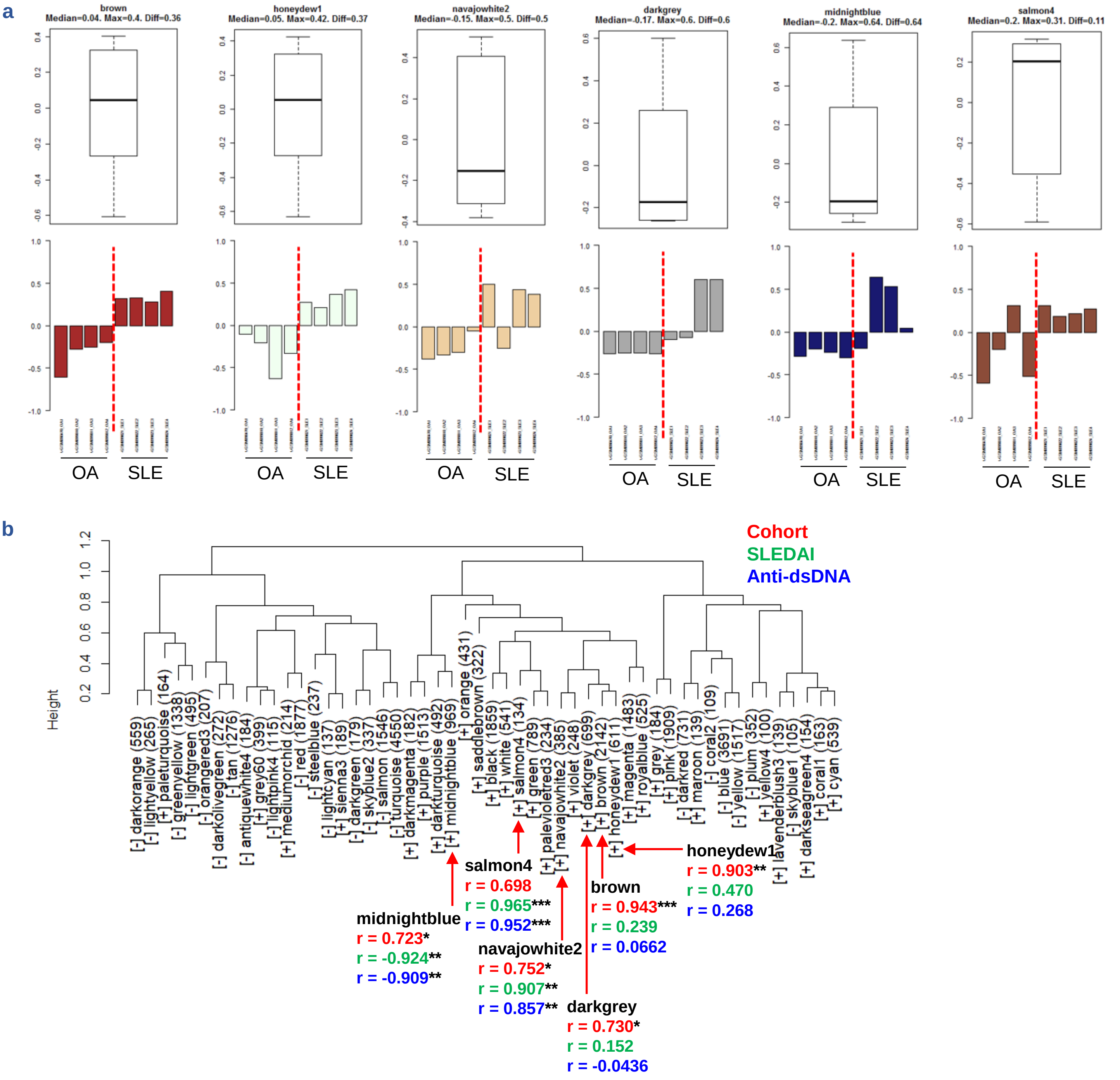


WGCNA of SLE vs OA patients yielded six modules of genes associated with LA after quality control. **(a)** Module eigengene plots per sample of the six LA-associated modules; color names are randomly generated as part of WGCNA module assignment. **(b)** Dendrogram of all 52 modules of co-expressed genes from WGCNA. Modules are labeled with the correlation direction to LA in brackets, the randomly generated color name of the module, and the number of microarray probes contained in the module in parentheses ([correlation] module name (# probes)). The six modules associated with LA are indicated with red arrows. Pearson correlation values to cohort, SLEDAI, and anti-dsDNA titer are shown in red, green, and blue, respectively. Significant correlations are marked: *p<0.05, **p<0.01, ***p<0.001.

### Supplementary Figure S2. Quality Control of WGCNA Results

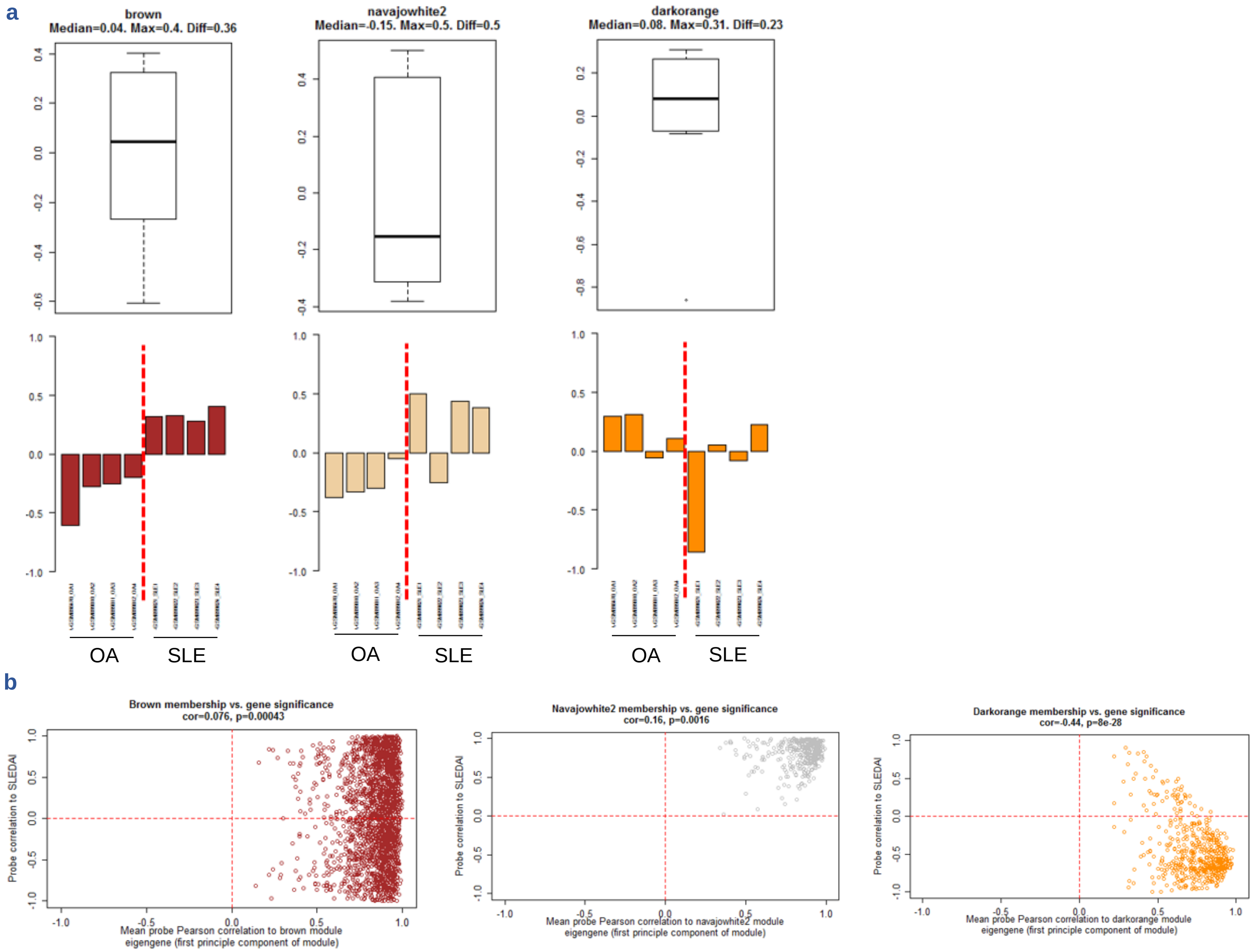


Two of the three WGCNA outputs contributing to quality control for the selection of modules for further interrogation are summarized here with three sample modules: brown, navajowhite2, and darkorange. **(a)** Module eigengene bar plots per patient sample in a module show variance of gene expression within a cohort. **(b)** Module membership (kME) plotted against individual probe correlation to SLEDAI shows agreement of individual module probes to trait correlation.

### Supplementary Table S1. Sample WGCNA Module Eigengene Correlations

| Module | cohort | | SLEDAI | | Anti-dsDNA | | C3 | | C4 | | CRP | |
|---|---|---|---|---|---|---|---|---|---|---|---|---|
| | r | p | r | p | r | p | r | p | r | p | r | p |
| brown | 0.943 | 4.3e-4 | 0.239 | 0.569 | 0.0662 | 0.876 | -0.0914 | 0.830 | 0.375 | 0.360 | -0.437 | 0.279 |
| navajowhite2 | 0.752 | 0.0315 | 0.907 | 0.00186 | 0.857 | 0.00653 | -0.928 | 8.90e-4 | -0.314 | 0.448 | 0.714 | 0.0468 |
| darkorange | -0.468 | 0.243 | -0.582 | 0.130 | -0.758 | 0.0294 | 0.0771 | 0.856 | -0.401 | 0.325 | -0.0644 | 0.880 |

Pearson correlations to clinical metadata, the final of three WGCNA outputs contributing to quality control for the selection of modules for further interrogation, are summarized here with three sample modules: brown, navajowhite2, and darkorange. Pearson correlation coefficients with corresponding p-values are described. Red and blue text indicate r values of significant correlations and anticorrelations ($p<0.05$), respectively.

**Supplementary Figure S3. Immune Cell Enrichment in LA-Associated WGCNA Modules**

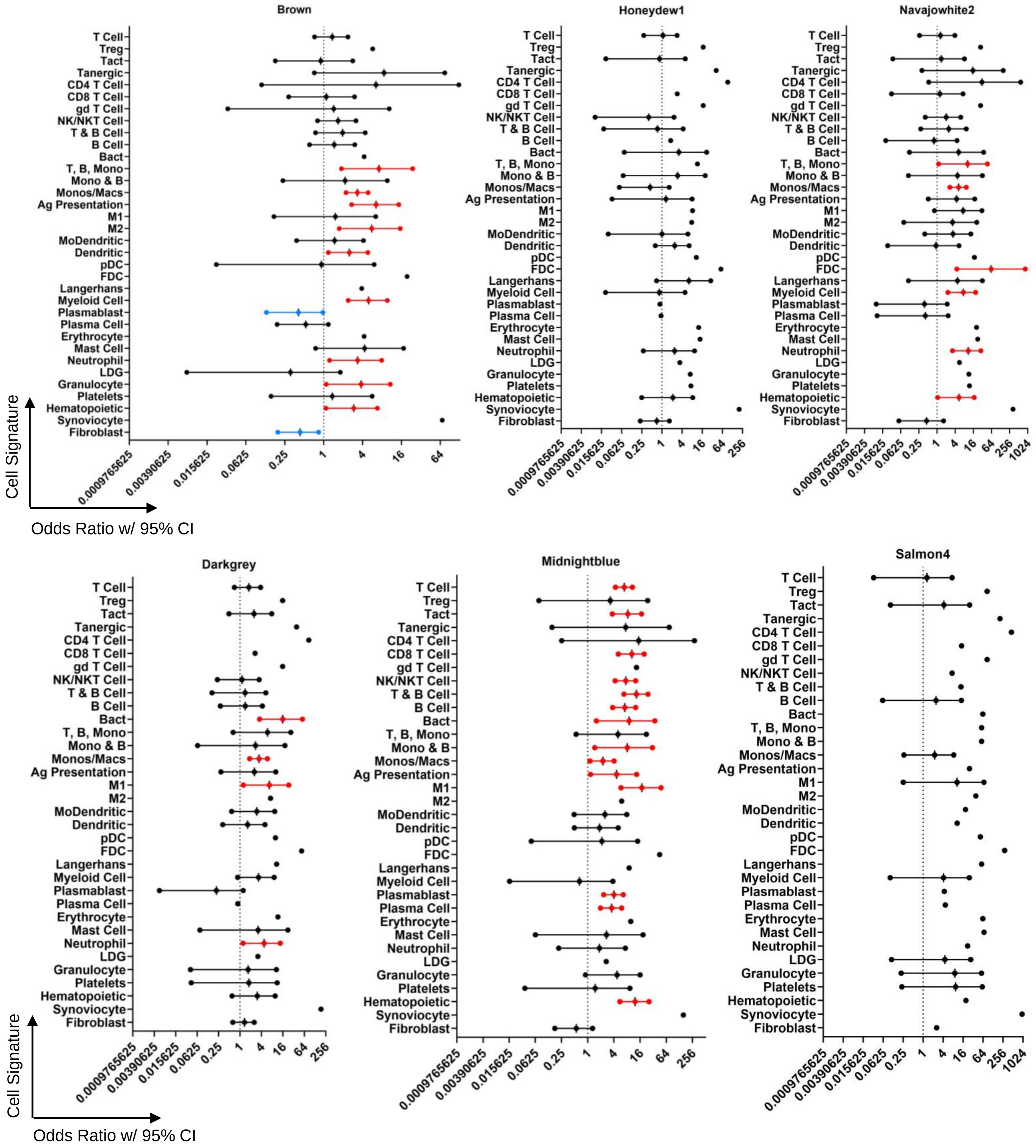


Odds ratios bound by 95% confidence intervals show specific immune/inflammatory cell populations or synovium-specific cell populations that could be linked to lupus synovitis. Significant enrichment ($p<0.05$) and confidence intervals that exclude odds ratio = 1 are colored red and blue for positive and negative association with the sample, respectively. X-axes are plotted on a log2 scale. For categories represented by a single point, odds ratio = 0 and the data point shown represents the upper bound of the confidence interval.

**Supplementary Figure S4. Characterization of WGCNA Modules Negatively Correlated with LA**

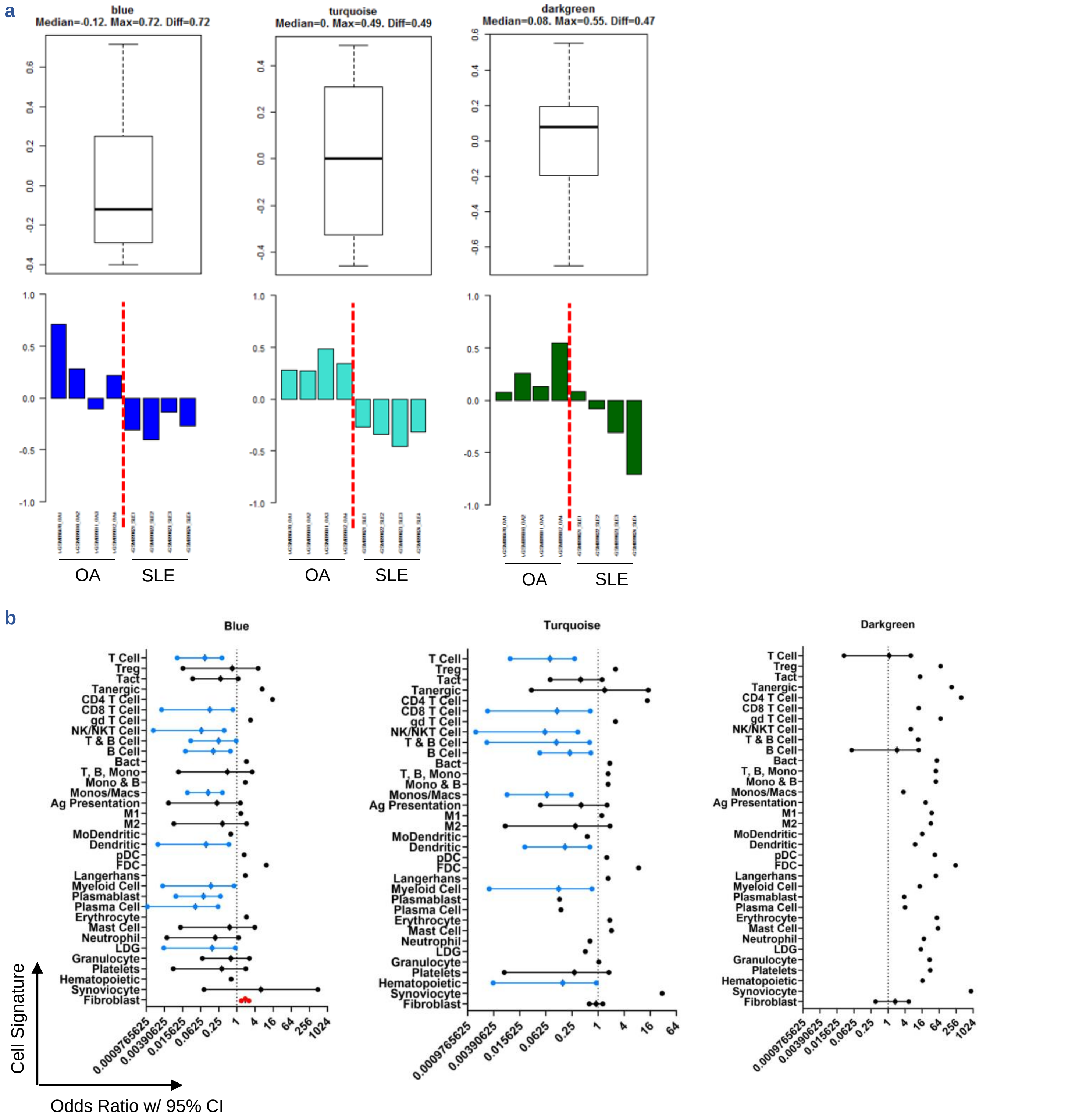


Overview of three WGCNA modules (blue, turquoise, darkgreen) negatively correlated with SLE. **(a)** Module eigengene bar plots per patient per cohort. **(b)** Enrichment of specific immune/inflammatory and synovial tissue cell populations. Significant enrichment ($p<0.05$) and a 95% confidence interval that excludes odds ratio = 1 are colored red and blue for positive and negative association with the sample, respectively. The x-axes are plotted on log2 scales. For categories represented by a single point, odds ratio = 0 and the data point shown represents the upper bound of the confidence interval.

**Supplementary Figure S5. Immunoglobulin Transcripts Identified in LA**

**a**

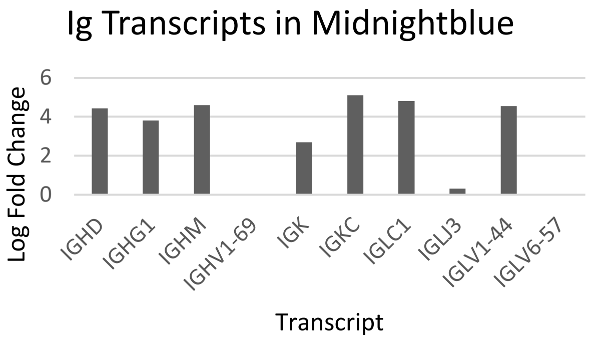


**b**

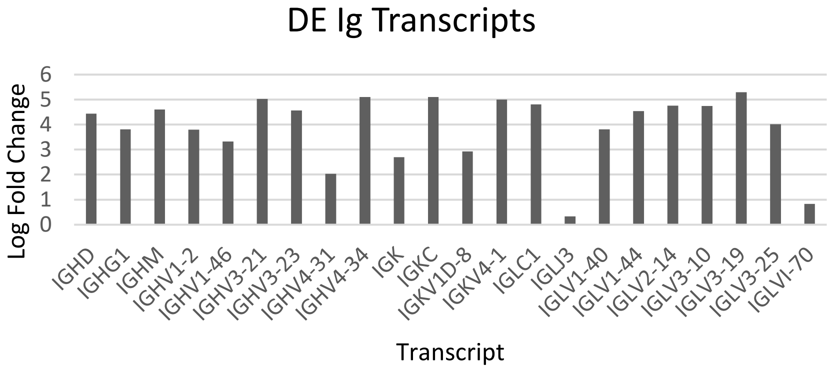


Immunoglobulin (Ig) transcripts detected in SLE vs OA synovium. **(a)** Log fold changes of Ig transcripts found in the LA-correlated WGCNA module containing the plasmablast/plasma cell signature, midnightblue. **(b)** Log fold changes of all Ig transcripts identified by differential expression analysis.

### Supplementary Table S2. WGCNA Modules Negatively Correlated with LA

| Module | cohort | | SLEDAI | | Anti-dsDNA | | C3 | | C4 | | CRP | |
|---|---|---|---|---|---|---|---|---|---|---|---|---|
| | r | p | r | p | r | p | r | p | r | p | r | p |
| blue | -0.791 | 0.0195 | 0.344 | 0.404 | 0.261 | 0.533 | -0.865 | 0.00557 | -0.894 | 0.00274 | 0.985 | 1.00e-5 |
| turquoise | -0.975 | 4.00e-5 | 0.395 | 0.333 | 0.430 | 0.288 | 0.284 | 0.495 | 0.942 | 4.70e-4 | -0.717 | 0.0454 |
| darkgreen | -0.716 | 0.0460 | -0.0778 | 0.855 | 0.159 | 0.707 | 0.509 | 0.198 | 0.449 | 0.264 | -0.247 | 0.555 |

Pearson correlation coefficients with corresponding p-values of all WGCNA modules significantly anticorrelated with LA. Red and blue text indicate r values of significant ($p<0.05$) correlations and negative correlations, respectively.

**Supplementary Figure S6. Functional Enrichment in LA-Associated WGCNA Modules**

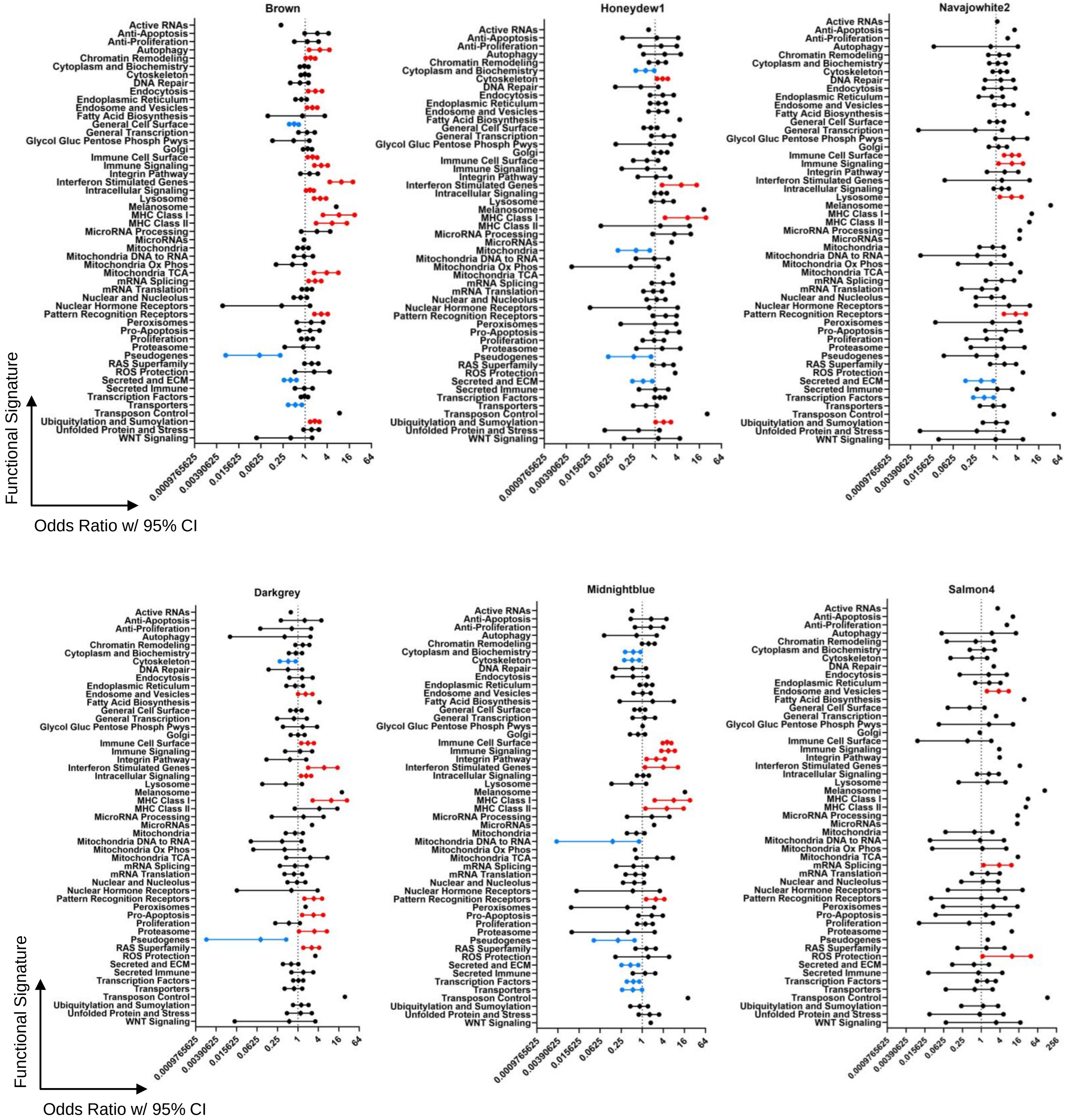


Odds ratios bound by 95% confidence intervals indicate enrichment of functional gene categories in six LA-associated WGCNA modules. Significant enrichment ($p<0.05$) and confidence intervals that exclude odds ratio = 1 are colored red and blue for positive and negative association with the sample, respectively. X-axes are plotted on a log2 scale. For categories represented by a single point, odds ratio = 0 and the data point shown represents the upper bound of the confidence interval.

### Supplementary Figure S7. Functional Characterization of LA-Associated WGCNA Modules

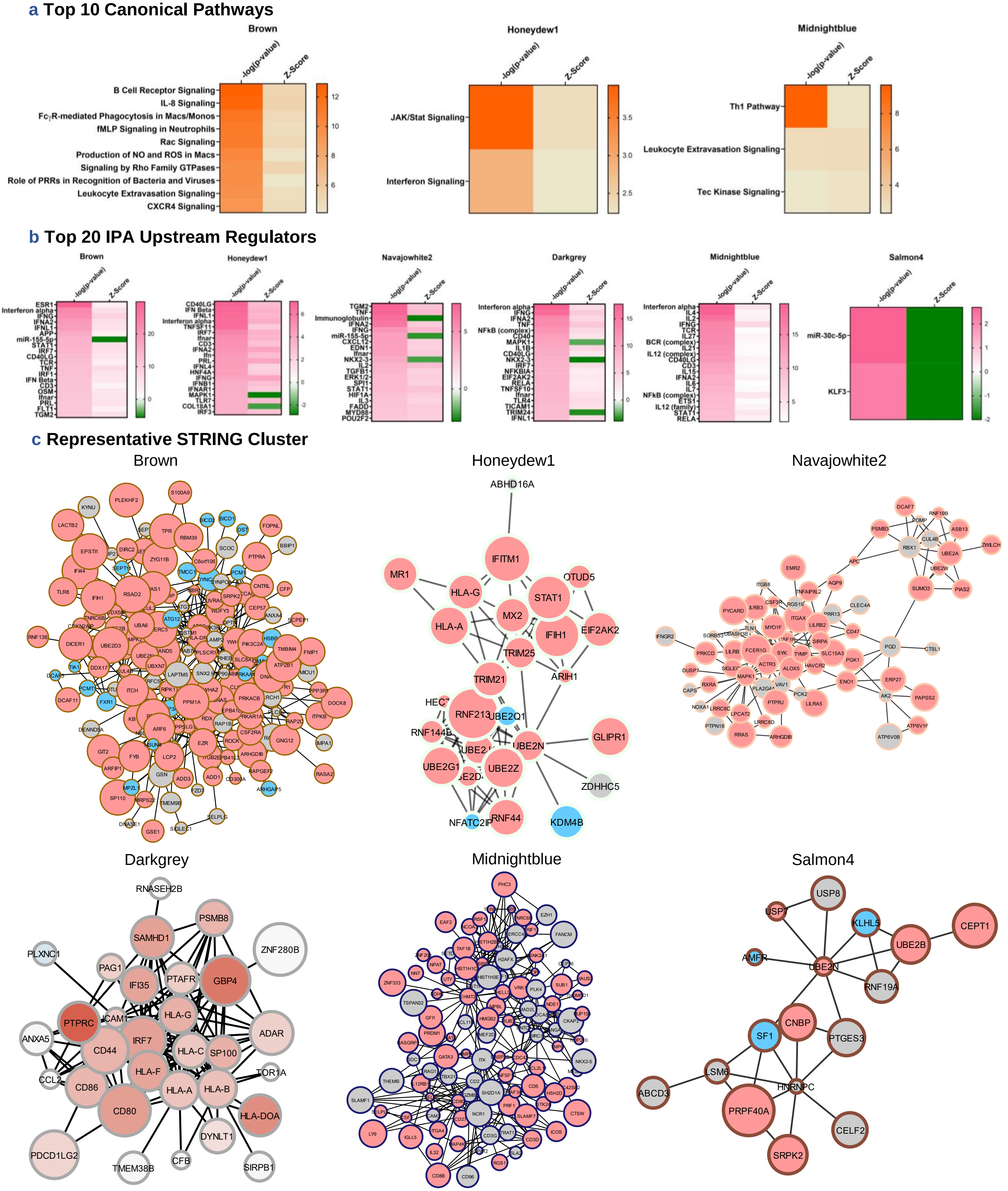


**(a)** Top 10 IPA canonical pathways for each of the six LA-associated WGCNA modules. Modules not shown were not predicted to have any significant pathways by p-value (p<0.01) and Activation Z-Score (|Z-Score| ≥ 2) with biological relevance. **(b)** Top 20 IPA upstream regulators for each LA-associated WGCNA module. All IPA predictions in **(a, b)** are ordered by significance (p<0.01) and must have an |Activation Z-Score| ≥ 2. **(c)** Single representative clusters from STRING depicting the protein-protein interactome of each LA-associated module. Clusters were chosen based on the highest number of intra-cluster connections and/or biological significance.

**Supplementary Figure S8. Germinal Center B Cell and $T_{fh}$ Cell Markers in LA**

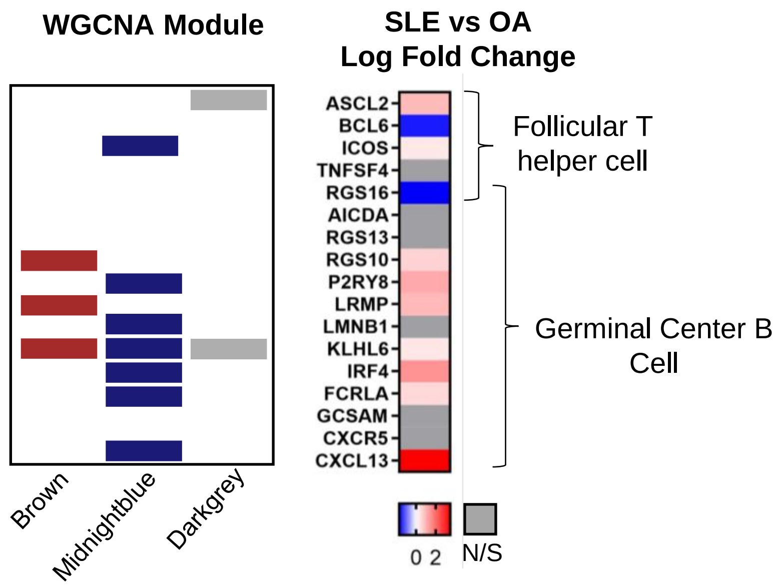


Assessment of GC B cell and $T_{fh}$ cell markers in lupus synovium from differential expression analysis and WGCNA. Genes found in SLE-associated WGCNA modules are indicated. LIMMA log fold changes indicate markers that are DEGs.

**Supplementary Figure S9. Comparison of Tissue-Specific Cell Types and Processes in SLE and RA Synovitis**

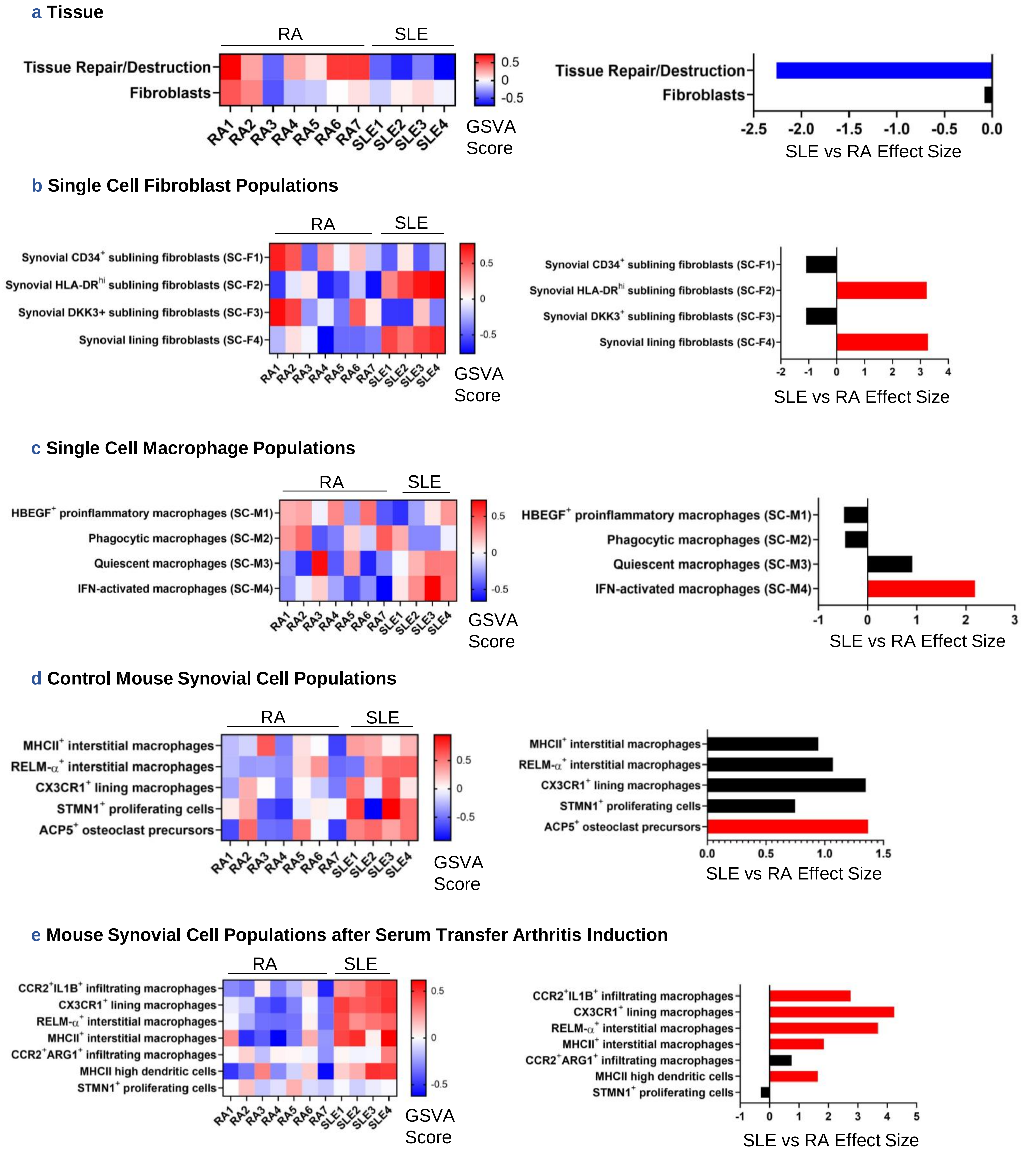


GSVA of synovial tissue processes and specific cell types **(a)** and literature-derived synovium-specific cell subtypes in human RA, OA, and mouse synovium **(b-e)** was conducted on log2-normalized gene expression values from SLE and RA synovium. Hedge's g effect sizes were calculated with correction for small sample size for each gene set and significant differences in enrichment between cohorts were found by Welch's t-test ($p<0.05$), shown in the panels on the right. Red and blue effect size bars represent significant enrichment in SLE and RA, respectively. Literature-derived signatures in **(b-e)** underwent co-expression analyses before being used as GSVA gene sets (see Methods).

**Supplementary Table S3. Compounds with Potential to Target LA**

| Target | Count | Range | Mean ± SEM | Top LINCS Drug | Representative Drug |
|---|---|---|---|---|---|
| PPAR | 4 | (-83.29)—(-98.18) | -91.18±3.18 | T-0070907[P] | Darglitazone‡ |
| PPAR agonist | 6 | (-83.72)—(-90.21) | -87.12±1.08 | Clofibrate† | Rosiglitazone†[1] |
| Acetylcholine receptor | 7 | (-75.36)—(-96.78) | -87.56±2.89 | Vinburnine† | Aclidinium† |
| Acetylcholine receptor agonist | 5 | (-81.38)—(-96.96) | -90.85±3.03 | VU-0365114-2 | Aceclidine† |
| Ca channel | 11 | (-76.29)—(-99.25) | -90.31±2.14 | Verapamil† | Amlodipine† |
| Ca channel activator | 2 | (-85.55)—(-89.40) | -87.47±1.93 | BAY-K8644[P] | FPL-64176[P] |
| Estrogen receptor | 3 | (-86.03)—(-91.93) | -89.30±1.73 | Fulvestrant† | Tamoxifen†[2] |
| Estrogen receptor agonist | 7 | (-78.36)—(-95.81) | -88.52±2.29 | PPT[P] | Dienestrol† |
| Dopamine receptor | 12 | (-76.84)—(-94.67) | -84.20±1.58 | Trifluoperazine† | Acepromazine† |
| Dopamine receptor agonist | 5 | (-78.47)—(-94.16) | -89.02±2.78 | Fenoldopam† | Bromocriptine† |
| SIRT | 2 | (-84.93)—(-92.45) | -88.69±3.76 | EX-527‡ | EX-527‡ |
| SIRT agonist | 2 | (-81.76)—(-82.14) | -81.95±0.19 | SRT-1720[P] | Curcumin†[7] |
| Serotonin receptor | 11 | (-76.84)—(-96.02) | -88.31±1.78 | Mirtazapine† | Agomelatine† |
| Serotonin receptor agonist | 9 | (-75.73)—(-90.74) | -81.91±1.88 | Ipsapirone‡ | Almotriptan† |
| GABA-A receptor | 4 | (-80.61)—(-93.42) | -86.65±3.23 | Flumazenil† | Butalbital† |
| GABA-A receptor agonist | 5 | (-81.76)—(-94.57) | -88.25±2.34 | Androstenol | Bemegride† |
| K channel | 4 | (-77.77)—(-96.79) | -86.59±4.03 | Linopirdine‡ | Amifampridine† |
| K channel activator | 2 | (-78.17)—(-79.89) | -79.03±0.86 | BMS-191011[P] | Diazoxide† |
| Progesterone receptor | 2 | (-81.77)—(-86.28) | -84.03±2.26 | Gestrinone† | Gestrinone† |
| Progesterone receptor agonist | 4 | (-76.92)—(-94.67) | -86.16±3.82 | Altrenogest† | Progesterone† |
| Adrenergic receptor | 13 | (-75.99)—(-96.02) | -86.15±1.81 | Mirtazapine† | Acebutolol† |
| Adrenergic receptor agonist | 6 | (-77.71)—(-94.43) | -84.44±2.47 | Dobutamine† | Epinephrine† |
| Glutamate receptor | 4 | (-81.61)—(-90.89) | -85.48±2.22 | Dextromethorphan† | Memantine†[0] |
| Glutamate receptor agonist | 2 | (-79.16)—(-86.30) | -82.73±3.57 | PHCCC[P] | Aniracetam† |
| Histamine receptor | 11 | (-75.02)—(-94.14) | -84.60±1.97 | Trimethobenzamide† | Acrivastine† |
| Histamine receptor agonist | 3 | (-77.56)—(-86.09) | -80.43±2.83 | Lidocaine† | Histamine† |

Select LINCS compounds for which both inhibitors/antagonists and activators/agonists were simultaneously predicted as targeting lupus synovitis. Compounds were analyzed if corresponding connectivity scores fell in the range of [-75 to -100] to reflect the most opposite gene signatures. Top LINCS Drug represents the most negative-scoring compound for a specific target category. Representative Drug conveys the most immunologically relevant or well-known drug for a specific target category. Where applicable, CoLTS scores [41] (range -16 to +11) are displayed as integers in superscript.

[P]: Preclinical

‡: Drug in development/clinical trials

†: FDA-approved

### Supplementary Figure S10. Co-expression Analysis of Synovial Cell Populations in SLE and OA Samples

#### a Co-expression of Single Cell Fibroblasts in SLE and OA

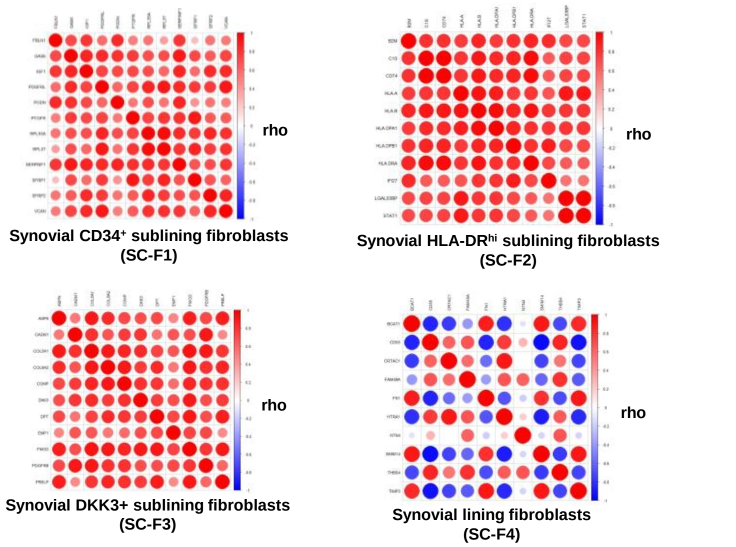


#### b Co-expression of Single Cell Macrophages in SLE and OA

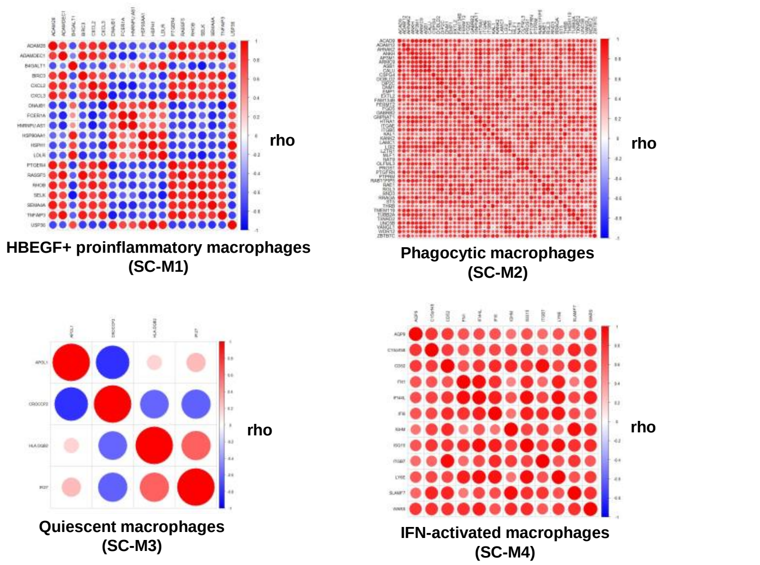


#### c Co-expression of Control Mouse Synovial Cell Populations in SLE and OA

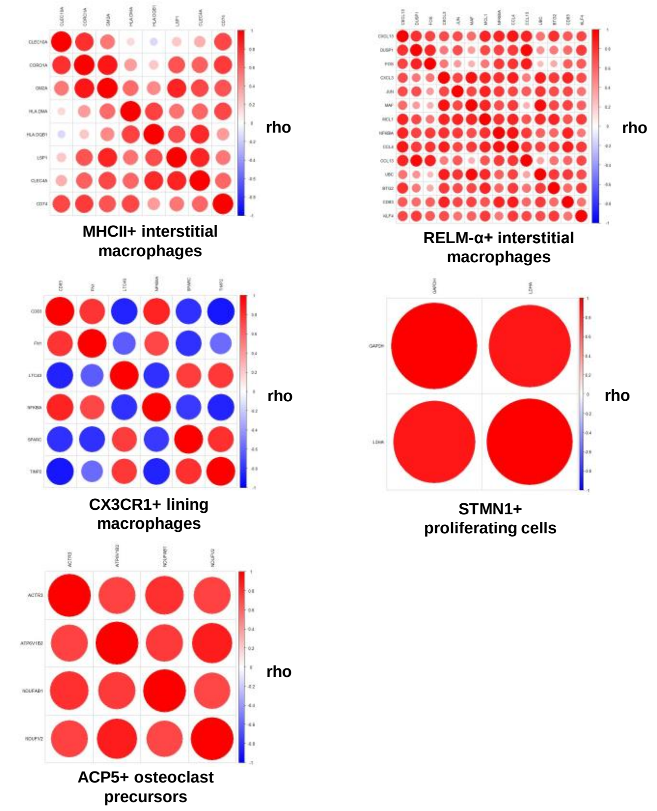


#### d Co-expression of Mouse Synovial Cell Populations after Serum Transfer Arthritis Induction in SLE and OA

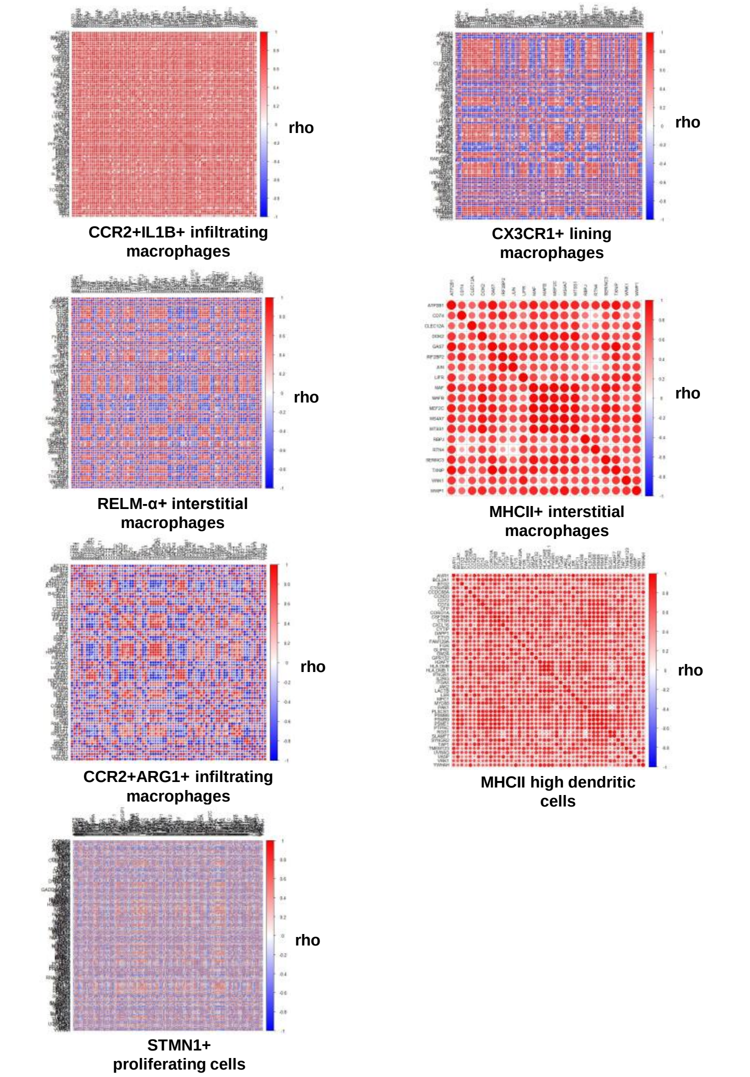


Spearman rank correlation coefficients (rho) of gene transcripts comprising synovial tissue-specific cell signatures computed from log2 expression values in SLE and OA samples. Refined signatures shown were derived from single-cell RNA-seq analysis of synovial fibroblasts **(a)** and macrophages **(b)** [10-11], normal mouse synovial cells **(c)** [12], and mouse synovial cells after serum transfer arthritis induction **(d)** [12]. Original signatures were culled based on percentage of significant correlations and Spearman rank rho values (see Methods).
